# Evolution of Yin and Yang isoforms of a chromatin remodeling subunit results in the creation of two genes

**DOI:** 10.1101/616995

**Authors:** Wen Xu, Lijiang Long, Yuehui Zhao, Lewis Stevens, Ronald E. Ellis, Patrick T. McGrath

## Abstract

Genes can encode multiple isoforms, broadening their functions and providing a molecular substrate to evolve phenotypic diversity. Evolution of isoform function is a potential route to adapt to new environments. Here we show that *de novo*, beneficial alleles in the *nurf-1* gene fixed in two laboratory strains of *C. elegans* after isolation from the wild in 1951, before methods of cryopreservation were developed. *nurf-1* encodes an ortholog of BPTF, a large (>300kD) multidomain subunit of the NURF chromatin remodeling complex. Using CRISPR-Cas9 genome editing and transgenic rescue, we demonstrate that in *C. elegans*, *nurf-1* has split into two, largely non-overlapping isoforms (NURF-1.B and NURF-1.D, which we call Yin and Yang) that share only two of 26 exons. Both isoforms are essential for normal gametogenesis but have opposite effects on male/female gamete differentiation. Reproduction in hermaphrodites, which involves production of both sperm and oocytes, requires a balance of these opposing Yin and Yang isoforms. Transgenic rescue and genetic position of the fixed mutations suggest that different isoforms are modified in each laboratory strain. In a related clade of *Caenorhabditis* nematodes, the shared exons have duplicated, resulting in the split of the Yin and Yang isoforms into separate genes, each containing approximately 200 amino acids of duplicated sequence that has undergone accelerated protein evolution following the duplication. Associated with this duplication event is the loss of two additional *nurf-1* transcripts, including the long-form transcript and a newly identified, highly expressed transcript encoded by the duplicated exons. We propose these lost transcripts are non-functional biproducts necessary to transcribe the Yin and Yang transcripts in the same cells. Our work suggests that evolution of *nurf-1* isoforms in nematodes creates adaptive conflict that can be resolved by the creation of new, independent genes.

## Introduction

There is general interest in understanding how animals adapt to new environments. What are the alleles that matter to positive selection and what sort of genes do they target? Since methods were developed to map and identify the genes harboring causative genetic variation, researchers have often isolated changes in the same gene in different populations or species (Wood, Burke et al. 2005, Martin and Orgogozo 2013). For example, in high-throughput experimental evolution studies in yeast, 77 putatively adaptive mutants occurred in just six genes (Venkataram, Dunn et al. 2016). Parallel evolution also seems to occur in multicellular organisms. Studies of pigmentation differences have repeatedly mapped genetic variation to the *agouti* and the *MC1R* genes (Flanagan, Healy et al. 2000, Hoekstra and Nachman 2003, Nachman, Hoekstra et al. 2003, Steiner, Weber et al. 2007, Linnen, Kingsley et al. 2009, Jones, Mills et al. 2018).

Besides targeting specific genes, evolution can target classes of genes that share molecular features such as biochemical (*e.g.* chemoreceptor genes (Bachmanov and Beauchamp 2007, Keller, Zhuang et al. 2007, Wisotsky, Medina et al. 2011, Lunde, Egelandsdal et al. 2012, McRae, Mainland et al. 2012, McBride, Baier et al. 2014, Greene, Brown et al. 2016, Greene, Dobosiewicz et al. 2016)) or developmental function (*e.g.* master regulators of cell fate (Sucena, Delon et al. 2003, Colosimo, Hosemann et al. 2005, Chan, Marks et al. 2010, Yang, Wang et al. 2018)). Another molecular feature predicted to be important for evolution is the ability of genes to produce multiple protein isoforms. A single protein-coding gene can produce multiple isoforms using alternative transcription initiation and termination sites combined with alternative splicing between exons (Pan, Shai et al. 2008, Pal, Gupta et al. 2011). Isoform-specific evolution is found throughout vertebrates, including recent evolution of transcript expression in primates (Barbosa-Morais, Irimia et al. 2012, Merkin, Russell et al. 2012, Shabalina, Ogurtsov et al. 2014, Zhang, Wang et al. 2017). Whether the increase in transcriptomic diversity is important for evolution remains an important question, and only a few examples have shown how isoform evolution could be involved in phenotypic diversity (Mallarino, Linden et al. 2017).

As a model for understanding the genetic basis of adaptive evolution in an animal model, we study two laboratory *Caenorhabditis elegans* strains, called N2 and LSJ2, which descended from a single hermaphrodite isolated in 1951 (**Figure 1A**). These two lineages split from genetically identical populations between 1957 and 1958 and evolved in two very different laboratory environments – N2 grew on agar plates seeded with *E. coli* bacteria and LSJ2 in liquid cultures containing liver and soy peptone extracts (McGrath, Rockman et al. 2009, McGrath, Xu et al. 2011, Sterken, Snoek et al. 2015). By the time permanent means of cryopreservation were developed, approximately 300-2000 generations had passed, and ∼300 new mutations arose and fixed in one of the two lineages (McGrath, Xu et al. 2011). Despite their genetic similarity, substantial divergence has occurred between these strains in terms of phenotype and evolutionary fitness, including a large number of developmental, behavioral, and reproductive traits. The low genetic diversity between these strains enables identification of not only the causal genes for these traits, but the exact causal nucleotides.

**Figure 1.**
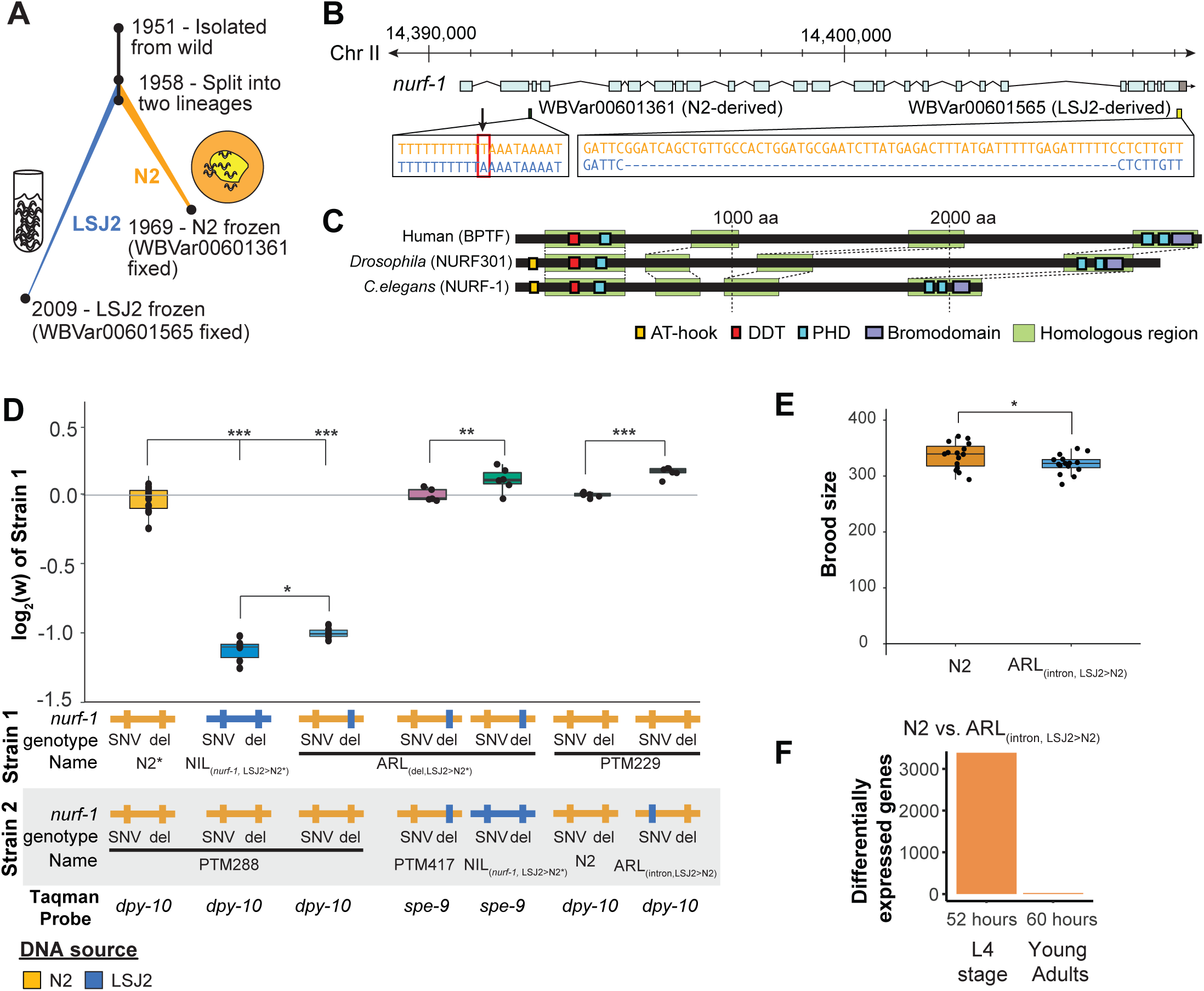
An N2-derived genetic variant in the intron of *nurf-1* increases fitness in laboratory conditions. **A**) History of two laboratory adapted *C. elegans* strains N2 and LSJ2, which descend from the same individual hermaphrodite isolated in 1951. The N2 and LSJ2 lineage split sometime around 1958. N2 grew on agar plates with *E.coli* OP50 as a food source for around 11 years until they were cryopreserved. LSJ2 animals were cultured in liquid axenic media containing sheep liver extract and soy extract peptone as a food source for about over 50 years until they were cryopreserved. 302 genetic variations were fixed between these two strains, including two that fall in the *nurf-1* gene – WBVar00601361 and WB00601565. **B**) Genetic location of two *nurf-1* variations. WBVar00601361 (in red box) is an N2-derived intron single nucleotide substitution T/A (N2/ancestral) in the 2nd intron of *nurf-1*. WBVar00601565 is an LSJ2-derived 60bp deletion in the 3’ end of *nurf-1* that removes the last 18 amino acids and part of the 3’-UTR. **C**) Comparison of NURF-1 orthologs from *Drosophila* and humans showing position of protein domains and conserved regions as determined by Blastp and Clustal Omega. **D**) Boxplot of pairwise evolutionary fitness differences between the indicated strains measured by directly competing the indicated strains against each other for five generations. PTM288 and PTM229 are the same genotype as N2* and N2, respectively, with the exception of an engineered DNA barcode in the *dpy-10* gene. PTM88 is the same genotype as the ARL_(del_LSJ2>N2)_, with the exception of a background DNA barcode in the *spe-9* gene (for details see **Methods**). The genotype of each *nurf-1* allele (shown in **B**) is indicated by color. The NIL strain also contains LSJ2 alleles of additional linked mutations, which is indicated by the blue horizontal line. For all figures, each dot represents an independent replicate, the box indicates the interquartile values of all data, and the line indicates the median of all data. Positive values indicate strain one is more fit than strain two. Negative values indicate strain two is more fit than strain one. For all figures, n.s. indicates p>0.05, one star indicates significant difference at p<0.05 level, two stars indicate significant difference at p<0.01 level, and three stars indicate significant difference at p<0.001 level. **E**) Total brood size of the N2 and ARL_(intron,LSJ2>N2)_ strains. **F**) Number of differentially expressed genes between synchronized N2 and ARL_(intron,LSJ2>N2)_ animals harvested 52 hours (L4 stage - when spermatogenesis is active) or 60 hours (young adults - when oogenesis is active) after hatching.

To date, five *de novo,* causal genetic variants have been identified in either the N2 or LSJ2 lineage (de Bono and Bargmann 1998, McGrath, Rockman et al. 2009, Persson, Gross et al. 2009, McGrath, Xu et al. 2011, Duveau and Felix 2012, Large, Xu et al. 2016, Large, Padmanabhan et al. 2017, Zhao, Long et al. 2018). Each of the derived alleles increases the evolutionary fitness of animals in the conditions they arose in. One of these mutations is an LSJ2-derived, 60 bp deletion at the 3’ end the *nurf-1* gene that reduces growth rate, slows reproductive output, and prevents development into the dauer diapause state in response to ascaroside pheromones (**Figure 1B**) (Large, Xu et al. 2016). This genetic variant is beneficial in the LSJ2 liquid cultures in which it arose and fixed, but places animals at a disadvantage in the agar plate environments in which N2 evolved, an example of gene-environment interaction (Large, Xu et al. 2016). We proposed that *nurf-1* is a regulator of life-history tradeoffs. Life history tradeoffs represent competing biological traits requiring large energetic investments, such as the tradeoff between energy required for reproduction versus the energy required for individual survival. The difference in fitness of this allele in the two laboratory environments is potentially determined by how the life-history tradeoffs map into reproductive success.

Studies of *nurf-1* and its orthologs provide fundamental support for its role as a life history regulator. *nurf-1* encodes an ortholog of mammalian BPTF, a subunit of the NURF chromatin remodeling complex (Barak, Lazzaro et al. 2003) (**Figure 1C**). *BPTF* encodes a large protein containing a number of domains that facilitate recruitment of NURF to specific regions of the genome for chromatin remodeling (Alkhatib and Landry 2011), including domains that interact with sequence-specific transcription factors and three PHDs and a bromodomain that facilitate interactions with modified nucleosomes (Li, Ilin et al. 2006, Wysocka, Swigut et al. 2006, Kwon, Xiao et al. 2009, Ruthenburg, Li et al. 2011). Through its DDT domain (Fyodorov and Kadonaga 2002), BPTF cooperates with ISWI to slide nucleosomes along DNA, changing access of promoter regions to transcription factors that drive gene transcription. In mammals, BPTF regulates cellular differentiation and homeostasis of specific cell-types and tissues, including the distal visceral endoderm (Landry, Sharov et al. 2008), ecoplacental cone (Goller, Vauti et al. 2008), hematopoietic stem/progenitor cells (Xu, Cai et al. 2018), mammary stem cells (Frey, Chaudhry et al. 2017), T-cells (Wu, Wang et al. 2016), and melanocytes (Koludrovic, Laurette et al. 2015). In *Drosophila*, the ortholog to BPTF, NURF301, regulates the heat shock response, pupation, spermatogenesis, and innate immunity (Badenhorst, Voas et al. 2002, Badenhorst, Xiao et al. 2005, Kwon, Xiao et al. 2008, Kwon, Xiao et al. 2009). Many of these traits can be viewed as life-history tradeoffs, *e.g.* large energetic investments in individual survival through the development of the immune system vs. energetic transfers to offspring in the placenta or mammary glands.

The evolution of BPTF/NURF-1 function might also be relevant in certain contexts. Genetic alterations of BPTF has an increasingly studied role in certain cancer progressions, and can be used as a diagnostic tool to predict patient outcomes (Xiao, Kim et al. 2014, Dai, Lu et al. 2015, Dar, Nosrati et al. 2015, Xiao, Liu et al. 2015, Xiao, Liu et al. 2015, Dar, Majid et al. 2016, Lee, Kim et al. 2016, Shiraishi, Okada et al. 2016, Gong, Liu et al. 2017, Sant, Tao et al. 2017, Seow, Matsuo et al. 2017, Duan, Wang et al. 2018, Roussy, Bilodeau et al. 2018, Pan, Yuan et al. 2019, Zhang, Han et al. 2019, Zhao, Zheng et al. 2019). Alteration in BPTF/NURF-1 function is implicated in the recent evolution of dingoes (Zhang, Wang et al. 2018) and the nematode *C. briggsae* (Chen, Shen et al. 2014).

In this paper, we continue our studies of the evolution of the N2/LSJ2 laboratory strains. We demonstrate that an independent, beneficial mutation in the *nurf-1* gene was fixed in the N2 lineage, suggesting that *nurf-1* is a preferred genetic target for laboratory adaptation. To understand why *nurf-1* might be targeted, we explored the *in vivo* role in *C. elegans* development by taking advantage of CRISPR-Cas9 to test causal relationships that inform laboratory evolution and fitness effects. Our work suggests that the large, full-length isoform of *nurf-1*, primarily studied in mammals, is dispensable for development. Instead, two, largely non-overlapping isoforms are both essential for reproduction, having opposing effects on cellular differentiation of gametes into sperm or oocytes. Our results suggest that the ability of *nurf-1* to regulate life history tradeoffs is the result of exquisite regulation of NURF function through the balance of two competing isoforms, reminiscent of the principal of Yin and Yang. Finally, we present evidence that these two isoforms have split into separate genes in a clade of related nematodes, potentially resolving conflict between the Yin and Yang isoforms transcription and function.

### An N2-derived variant in the second intron of *nurf-1* increases fitness and brood size in laboratory conditions

We previously mapped differences in a number of traits (including reproductive rate, fecundity, toxin and anthelmintic sensitivity, and laboratory fitness) between N2 and LSJ2 to a QTL centered over *nurf-1,* which contains a derived mutation in both the N2 and LSJ2 lineages (**Figure 1A** and **B**) (Large, Xu et al. 2016, Large, Padmanabhan et al. 2017, Zhao, Wan et al. 2019). The LSJ2 allele of *nurf-1* contains a 60 bp deletion in the 3’ end of the coding region of the gene, overlapping the stop codon and probably resulting in the translation of parts of the 3’ UTR. The N2 allele of *nurf-1* contains an SNV that converts an A to a T in a homopolymer run of Ts in the 2^nd^ intron (**Figure 1B**). Using CRISPR-Cas9-based genome editing, we previously demonstrated that the LSJ2-derived deletion accounted for a large portion of the trait variance in reproductive rate explained by the QTL. However, it did not explain the entire effect of this locus (Large, Xu et al. 2016). We decided to test whether this additional genetic variant or variants affected evolutionary fitness of the animals in laboratory conditions using a previously described pairwise competition assay (Zhao, Long et al. 2018). To do so, we took advantage of three strains we had previously created; CX12311 is a near isogenic line used to eliminate the fitness and phenotypic effect of derived alleles of N2 *npr-1* and *glb-5* (Zhao, Long et al. 2018) (**Figure 1D** - referred to as N2*), NIL_(*nurf-1*,LSJ2>N2*)_ is a near isogenic line containing LSJ2 alleles of both *nurf-1* mutations backcrossed into an N2* background (Large, Xu et al. 2016), and ARL_(del,LSJ2>N2*)_ is an allelic replacement line containing the LSJ2-derived 60 bp deletion edited into the N2* strain using CRISPR-Cas9. Phenotypic differences between the NIL_(*nurf-1*,LSJ2>N2*)_ and ARL_(del,LSJ2>N2*)_ strains are caused by the N2-derived intron SNV in *nurf-1*, or one of the additional seven linked LSJ2-N2 genetic variants near *nurf-1*.

We measured the relative fitness of the N2*, NIL_(*nurf-1*,LSJ2>N2*)_, and ARL_(del,LSJ2>N2*)_ strains against PTM288, a version of N2* that also contains a silent mutation in the *dpy-10* gene (**Figure 1D**). The *dpy-10* silent mutation provides a common genetic variant that can be used to quantify the relative proportion of each strain on a plate using digital droplet PCR. Both the NIL_(*nurf-1*,LSJ2>N2*)_ and ARL_(del,LSJ2>N2*)_ strains showed dramatically reduced fitness comparing to PTM288, consistent with our previous report showing that the 60 bp deletion is deleterious on agar plates (Large, Xu et al. 2016). However, the NIL_(*nurf-1*,LSJ2>N2*)_ was quantitatively and significantly less fit than the ARL_(del,LSJ2>N2*)_ strain, suggesting additional genetic variant(s) in the NIL_(*nurf-1*,LSJ2>N2*)_ strain further reduced its fitness. To confirm this result, we also directly competed the NIL_(*nurf-1*,LSJ2>N2*)_ and ARL_(del,LSJ2>N2*)_ strains against each other, using a nearly neutral background mutation in *spe-9(kah132)* to distinguish the two strains (**Figure 1D**).

To determine if the N2-derived intron SNV in *nurf-1* (**Figure 1B**) was responsible for the fitness gains (as opposed to one of the seven linked LSJ2/N2 variants), we used CRISPR-Cas9 to directly edit the LSJ2 allele of the intron SNV into the standard N2 strain to create a strain we will refer to as ARL_(intron,LSJ2>N2)_. We measured the relative fitness of the ARL_(intron,LSJ2>N2)_ and N2 strains against PTM229 (a strain which again contains a *dpy-10* silent mutation). The ARL_(intron,LSJ2>N2*)_ strain was significantly less fit than the N2 strain at a level similar to the difference between the NIL_(*nurf-1*,LSJ2>N2*)_ and ARL_(del,LSJ2>N2)_ strains (**Figure 1D)**. These results indicate that beneficial alleles of *nurf-1* arose in both laboratory lineages - the 60 bp deletion makes LSJ2 animals more fit in liquid, axenic media (Large, Xu et al. 2016), and the intron SNV makes N2 animals more fit on agar plates seeded with bacteria.

Brood size of *C. elegans* hermaphrodites is an important trait for evolutionary fitness in laboratory conditions and an example of a life-history tradeoff (Cutter 2004, Anderson, Reynolds et al. 2011, Murray and Cutter 2011). After sexual maturation, gonads in the hermaphroditic sex initially undergo spermatogenesis before transitioning to oogenesis; a concomitant lengthening of spermatogenesis time increases the total brood size of hermaphrodites but also delays when reproduction can start. When we compared the total fecundity produced by the N2 and ARL_(intron,LSJ2>N2)_ strains, we found a significant difference, with the ARL_(intron,LSJ2>N2)_ strain producing ∼30 fewer offspring than N2 (**Figure 1E**). The reproductive rate of the N2 and ARL_(intron,LSJ2>N2)_ strains was largely unchanged throughout their reproductive lifespan (**Figure S1**).

RNAseq analysis identified transcriptional differences caused by the intron SNV during spermatogenesis, supporting our hypothesis that sperm development is affected by this SNV. We collected RNA from synchronized N2 and ARL_(intron,LSJ2>N2)_ hermaphrodites at two timepoints, 52 and 60 hours after hatching, which occur during spermatogenesis (52 hours) or oogenesis (60 hours). Interestingly, a large number of genes are differentially expressed between the two strains but only during the 52-hour timepoint (3,384 genes vs. 25 genes) (**Figure 1F, Figure S2A**, and **Table S1**). Although a portion of these 3,384 genes are expressed in the germline, these genes are also expressed in additional tissues (**Figure S2B**). Gene ontology analysis suggests that cuticle development and innate immune responses are regulated by *nurf-1* (**Table S2**) consistent with the role of its orthologs in regulating immunity and melanocyte proliferation in *Drosophila* and humans (Kwon, Xiao et al. 2008, Landry, Banerjee et al. 2011, Koludrovic, Laurette et al. 2015, Wu, Wang et al. 2016). These results suggest that the intron SNV regulates a number of developmental processes including spermatogenesis, molting, and innate immunity.

### *nurf-1* produces multiple transcripts encoding multiple protein isoforms

Our results suggest that selection acted repeatedly on *C. elegans nurf-1* during laboratory growth. The molecular nature of NURF-1, an essential subunit of the NURF chromatin remodeling complex, is surprising for a hotspot gene. In general, chromatin remodelers are thought of as ubiquitously expressed regulators with little variation in different cell types, akin to general function RNA polymerase proteins or ribosomes. Why would genetic perturbation of *nurf-1* lead to increased fitness? One potential clue is the complexity of the *nurf-1* locus. Previous cDNA analysis of *nurf-1* identified four unique transcripts encoding four unique isoforms (Andersen, Lu et al. 2006), two of which have been shown to regulate different phenotypes (summarized in **Table 1**).

To identify other transcripts produced by *nurf-1* and quantify the relative proportions of each that are produced, we analyzed previously published Illumina short-read (Brunquell, Morris et al. 2016) (isolated from synchronized L2 larval animals) and Oxford Nanopore long-read RNA sequencing reads (Roach, Sadowski et al. 2019) (isolated from mixed populations) (**Figure S3, Figure S4**). Our results support many of the conclusions of Andersen *et al.* (Andersen, Lu et al. 2006) but contain a few surprises. We identified five major transcripts (**Figure 2A**) - three previously isolated (*nurf-1.b*, *nurf-1.d,* and *nurf-1.f*) but also two newly identified (*nurf-1.a* and *nurf-1.q*) (mapping of transcript names used in Andersen *et al.* are listed in **Table 1**). *nurf-1.a* encodes a full-length 2,197 amino acid isoform analogous to the primary isoform of BPTF in humans and NURF301 in *Drosophila* (**Figure 1C**). Despite the expectation that *C. elegans* would produce a similar protein, the Oxford Nanopore long-read data is the only evidence supporting its existence. The *nurf-1.q* transcript is predicted to produce a 243 amino acid unstructured protein. With the exception of the full-length *nurf-1.a* transcript, the overlap of these transcripts is quite minimal, resulting in predicted isoforms with unique protein domains and functions (**Figure 2B**).

**Figure 2.**
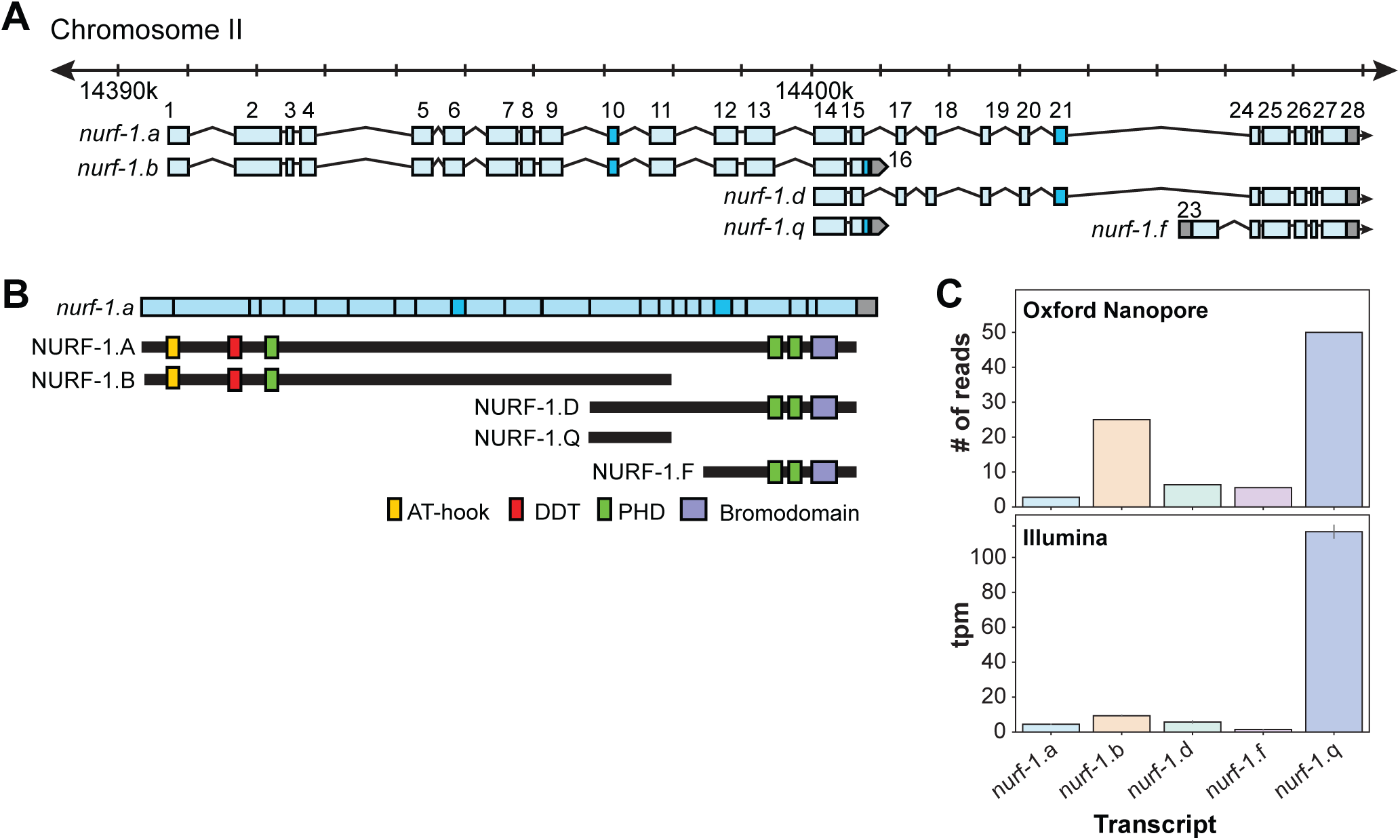
*nurf-1* encodes five major transcripts. **A**) Genomic position of the five *nurf-1* transcripts supported by Illumina short read and Oxford Nanopore long reads. Each blue box is an exon. Exon number is indicated on the figure. Dark blue exons (10, 16, and 21) are alternatively spliced, resulting in a 6-9 bp difference in length (see **Figure S4** for details). **B**) The predicted protein isoforms produced by each of the five major transcripts and along with the domains each isoform contains. Immunoblots only supported translation of the B, D, and F isoforms (see **Figure S5** for details). For reference, the spliced *nurf-1.a* transcript is also shown as reference. **C**) Relative expression levels of each transcript, determined by number of Oxford Nanopore reads from a mixed population (top panel) or analysis of Illumina short reads from L2 staged animals using kallisto (bottom reads). tpm = transcripts per million.

We quantified the relative expression of these five transcripts by either counting the number of Nanopore reads that matched the transcript or by using kallisto (Bray, Pimentel et al. 2016) to predict transcript abundance using Illumina short-read sequencing data (**Figure 2C**). These predictions qualitatively agreed in transcript ranking of expression strength (although quantitative variation in predictions were observed, reflective of the different technologies or developmental stages of the animals). Surprisingly, the newly described *nurf-1.q* transcript was the most highly expressed followed by the *nurf-1.b* transcript, and the *nurf-1.a, nurf-1.d* and *nurf-1.f* were expressed at similar lower levels.

Although each of the five major transcripts are transcribed, this result does not necessarily mean they are translated into stable protein products. To facilitate analysis of NURF-1 proteins, we used CRISPR-Cas9 to fuse two distinct epitope tags (HA and 3xFLAG tag) to the endogenous *nurf-1* locus, just prior to the stop codons in the 16^th^ and 28^th^ exon, respectively (**Figure S5A**). Immunoblot analysis supported the expression of the B, D, and F isoforms, but not the A or Q isoforms (**Figure S5B**). Although larger proteins, such as the A isoform, can be difficult to transfer during immunoblots, the lack of a band matching the small Q isoform suggests the highly expressed *nurf-1.q* transcript is not translated into protein or the protein is rapidly degraded.

### The B and D isoforms are both essential for reproduction and the F isoform modifies the heat shock response

Genetic analysis of *nurf-1* primarily relied on two deletion alleles, *n4293* and *n4295* (**Figure 3A**) (Andersen, Lu et al. 2006). The *n4293* allele deletes the first exon and predicted transcriptional start site of the *nurf-1.a* and *nurf-1.b* transcripts. The *n4295* allele deletes three exons of the *nurf-1.a*, *nurf-1.d*, and *nurf.1.f* transcripts that encode a C-terminal PHD domain (**Figure S6**) necessary for human BPTF function. Comparison of the phenotypes of the *n4293* and *n4295* homozygotes leads to the model that the B isoform is essential for reproduction and the A, D, and/or F isoforms have subtle effects on growth rate and reproductive rate (**Table 1**).

**Figure 3.**
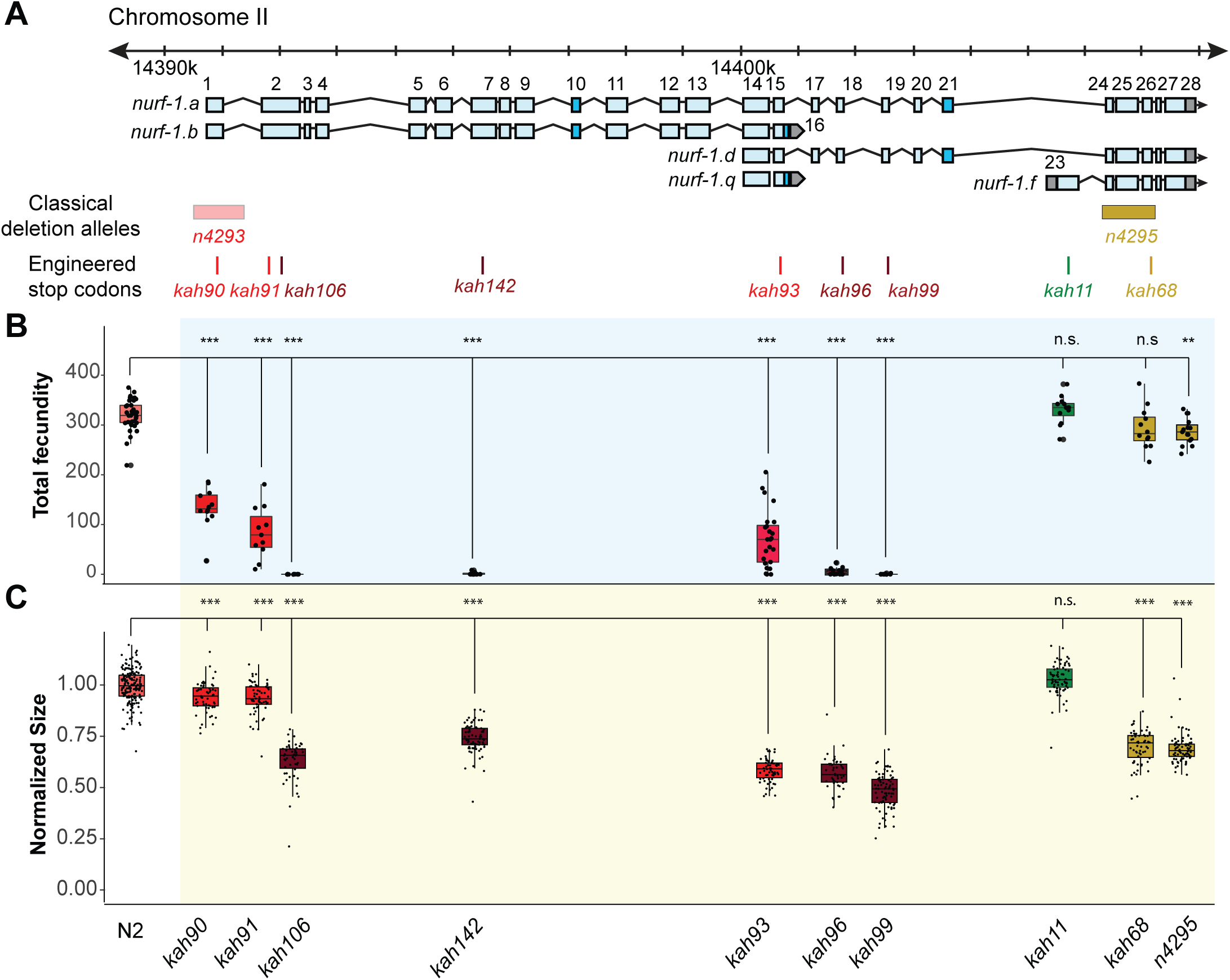
An additional isoform besides NURF-1.B is necessary for reproduction in *C. elegans*. **A**) Genomic positions of *nurf-1* classical deletion alleles and nine engineered stop codons created using CRISPR/Cas9 based gene editing. Each allele is color-coded by the reproductive ability of homozygous strains. Green is statistically indistinguishable from wild-type, yellow indicates slightly reduced brood size and change in reproductive rate, red indicates substantially reduced brood size in the first generation and eventual sterility after multiple generations of homozygosity, and dark red indicates sterility in the first generation of homozygosity. **B**) Fecundity of indicated strains (shown in x-axis of panel) **C**). For red or dark red strains (panel A), measurements were taken on animals homozygous for a single generation.

To further delineate the biological role of each isoform, we used CRISPR-Cas9 to engineer nine stop codons in eight exons of the *nurf-1* gene: the first, second (two positions), 7^th^, 15^th^, 18^th^, 19^th^, 23^rd^, or 26^th^ exons (**Figure 3A**). The predicted effects of these stop codons on each major isoform are shown in **Figure S6** and **Table 2**. Homozygote animals for each mutation were assayed for total brood size and growth rate. Analysis of the phenotypes of these mutants indicated that our working model was incorrect. Instead, we propose that both the B and D isoforms are essential for reproduction.

As expected, engineering stop codons in the first, second, and 7^th^ exons greatly reduced fecundity, resulting in either sterility, or a mortal germline phenotype, initially reducing total brood size of animals, before eventually causing complete sterility after around three-to-five generations of homozygosity (Figure 3B and C). Although the qualitative phenotypes of these four alleles agreed, we observed interesting quantitative differences between them. The second stop codon in the second exon (*kah106*) and the stop codon in the 7^th^ exon (*kah142*) reduced growth and fecundity more than the first exon stop codon (*kah90*) or the first stop codon in second exon (*kah91*) (Figure 3B and C). We believe this result indicates the presence of an internal ribosome entry site in the middle of the second exon at the 122^nd^ Methionine, causing the expression of two isoforms from a single transcript. The reduced severity of the first two stop codon alleles can be explained by their inability to affect the protein sequence of the second isoform. An alternative possibility is a difference in frequency of translational read-through of each stop codon, which are interpreted as sense codons at a low frequency (Jungreis, Lin et al. 2011).

Unexpectedly, engineering stop codons in the 18^th^ and 19^th^ exons also caused a mortal germline phenotype (*kah96* and *kah99*) (**Figure 3B**). This result was surprising, because the *n4295* allele, predicted to be a loss-of-function allele for the D and F isoforms due to the loss of the PHD and bromodomains, does not have a mortal germline phenotype. We excluded a number of potential explanations for this discrepancy. A suppressor for the *n4295* allele could have fixed during the construction of this strain. However, the *kah68* allele, which contains a stop codon within the *n4295* deleted region, phenocopies the *n4295* allele and not the *kah96* and *kah99* animals (**Figure 3B**, **3C**, and **Figure S7**)). Another possibility is that the D isoform suppresses the F isoform; loss of both isoforms (in the *n4295* background) is tolerated, but loss of just the D isoform (in the *kah96* or *kah99* backgrounds) allows the F isoform to prevent reproduction. However, we could exclude this possibility as the double mutant containing both the *n4295* allele and the 18^th^ exon stop allele phenocopied the *kah96* single mutant (**Figure S8**). Additionally, specific loss of the F isoform by the 23^rd^ exon stop allele (*kah11*) did not affect the phenotype of animals (Figure 3B and C). Our data suggests that, unlike human BPTF, the ability of NURF-1 to bind modified histones is not required for its function. We further confirmed this hypothesis by editing conserved residues in these the PHD and bromodomains necessary for recognition of the H3K4me3 and H4K16ac marks (**Figure S9**).

The most parsimonious explanation of our data is that either the A or D isoform is essential for reproduction in *C. elegans*. Compound heterozygote tests allowed us to distinguish between these possibilities, indicating that the D isoform is required for reproduction and wild-type growth rate, and the A isoform is dispensable for reproduction and development (**Figure 4**). We first verified that the *kah93*, *kah96*, and *kah106* alleles were recessive by measuring the fecundity of heterozygous animals (**Figure 4B**). Next, we examined the fecundity of *kah106/kah96* compound heterozygotes, which are predicted to lack only the A isoform, due to the production a single unaffected copy of the B isoform from the *kah96* haplotype and a single unaffected copy of the D isoform from the *kah106* haplotype. If the A isoform was essential for reproduction, we would expect these compound heterozygotes to be sterile or have severe defects in fecundity. However, these animals were indistinguishable from wild-type, suggesting that the full-length A isoform is not essential (**Figure 4B**). The *kah106/kah93* compound heterozygotes showed similar results. These animals are predicted to encode one unaffected copy of the D isoform, one truncated copy of the B isoform, and zero unaffected copies of the A isoform. These animals were mostly wild-type, with a small reduction in total fecundity (**Figure 4B**). We believe that the A isoform is not essential and the truncation of the B isoform slightly perturbs its function, causing a slight reduction in fecundity. Finally, we analyzed *kah93/kah96* compound heterozygotes. These animals are predicted to encode zero wild-type copies of the D isoform, one wild-type copy of the B isoform, and zero wild-type copies of the A isoform. These animals were essentially sterile. Taken together, we believe that the B and the D isoform are both essential for reproduction.

**Figure 4.**
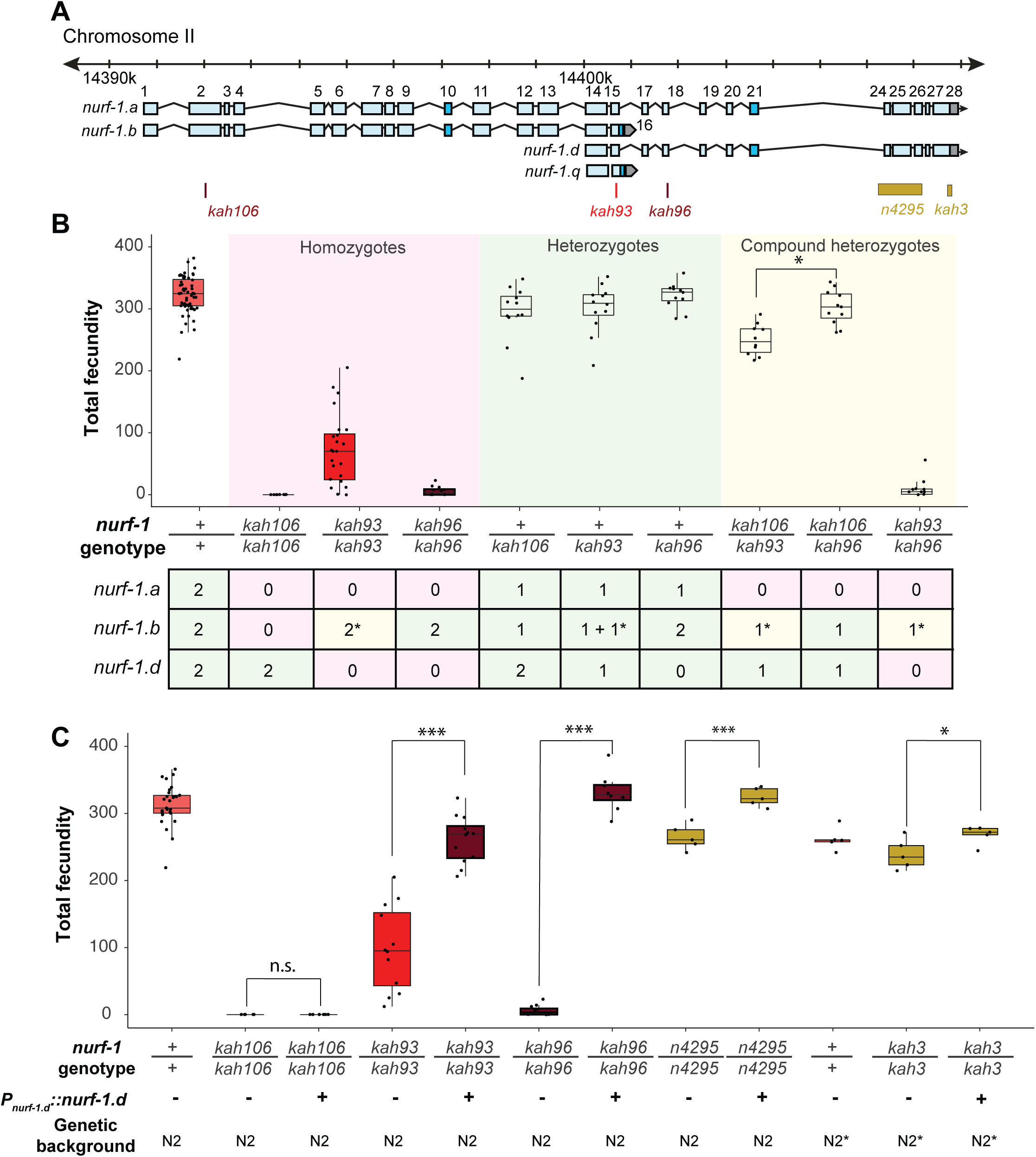
Genetic analysis suggests the NURF-1.B and NURF-1.D isoforms are essential for reproduction in *C. elegans*. **A**)Genomic positions of stop codon or classical deletion mutations used for compound heterozygote or transgenic rescue analysis of B and C. *kah3* is a CRISPR/Cas9 genomic edit of the LSJ2-derived 60 bp deletion. **B**) Fecundity of homozygote (red), heterozygote (green), and compound heterozygote mutants (yellow) as indicated in the x-axis. The table below the x-axis is the predicted effect of each mutant strain on the indicated *nurf-1* isoforms. The number in the table indicates the number of functional copies. The star indicates the milder predicted effect of *kah93* on NURF-1.B, as it only truncates 73 of 1,621 amino acids. The y-axis shows the fecundity for each strain. **C**) Fecundity of indicated strains with and without the presence of an integrated *nurf-1.d* transgene. The genetic background is also indicated. N2* contains ancestral introgressions of the *npr-1* and *glb-5* genes.

To confirm that the D isoform is essential, we also created a transgenic strain containing an integrated construct driving a *nurf-1.d* cDNA from its endogenous promoter. This transgene could fully rescue the fecundity phenotype of the *kah96* allele and partially rescue the fecundity phenotype of the *kah93* allele (**Figure 4C**). This transgene could also rescue the reproductive timing and fecundity changes of the *n4295* allele and the LSJ2-derived 60 bp deletion (*kah3*) (**Figure 4C** and **Figure S10**). As expected, this transgene could not rescue the *kah106* allele, which creates a stop codon in the B isoform. These data further support a requirement of both the B and D isoforms for reproduction.

Although the F isoform does not seem to have an effect in normally developing animals, it is involved in the heat shock response. Multiple reports have demonstrated that *nurf-1* is upregulated in response to heat shock (Brunquell, Morris et al. 2016, Li, Chauve et al. 2016). By analyzing these reads, we found that the *nurf-1.f* transcript was specifically upregulated in both datasets, with increased coverage of the 23^rd^ exon as well as the 24^th^ through 28^th^ exons (**Figure S11A and B**). We confirmed that the increased transcription of the *nurf-1.f* transcript also increased NURF-1.F protein abundance (Figure S11C and D). Transcriptional analysis of strains lacking the F isoform indicated that the initial transcriptional response to heat shock was largely the same, but the long-term transcriptional response of a subset of genes was affected (**Figure S11E-G**). We conclude that the F isoform is specifically up-regulated by heat shock and plays a modulatory role in determining the long-term transcriptional response to heat shock.

### The B and D isoforms have opposite effects on cell fate during gametogenesis

Although the B and D isoforms are both required for reproduction, the molecular mechanism that these isoforms operate through could be different. One possibility is that the long-form of NURF-1 has split into two subunits - both isoforms participate as part of the NURF complex, cooperating together to regulate reproduction. However, the D isoform might instead modify NURF activity by competing for binding with transcription factors or regions of the genome to which NURF is recruited. A third possibility is that the D isoform acts through a NURF-independent pathway.

To gain insights into the molecular nature of the D isoform, we decided to determine precisely how the B and D isoforms regulate reproduction, using three nurf-1 stop alleles (**Figure 5A**). For hermaphrodites to produce a fertilized egg, the gonads must produce both male and female gametes at different developmental times (**Figure 5B**). Initially, gametogenesis produces sperm, creating approximately 300 sperm at which point a permanent sperm-to-oocyte switch occurs. From this time, gametogenesis produces oocytes until the animal dies or the gonad ceases to function (Hubbard and Greenstein 2005). A number of defects could cause sterility – inability to form gametes, inability to create sperm, inability to create oocytes, or defects in the sperm and/or oocyte function. We used DAPI staining to characterize the production of sperm and oocytes in three *nurf-1* mutants (Figure 5C and D). We first tested *kah106* mutants, which lack the B isoform (**Figure 5A**), for the ability to produce sperm. Compared with N2 animals, which create ∼300 sperm per animal, the number of sperm produced by *kah106* animals was greatly reduced, resulting in the production of only approximately 60 sperm (**Figure 5D**). These animals produced a normal number of oocytes, indicating that spermatogenesis seemed to be affected specifically (**Figure 5E**). We interpret these data as evidence that hermaphrodites that lack the NURF-1.B isoform spend less time in spermatogenesis before transitioning to oogenesis. We next tested *kah96* mutants which lack the D isoform. These animals produced approximately 500 sperm (Figure 5C and D) and almost no oocytes (**Figure 5E**). We interpret these data as evidence that hermaphrodites that lack the D isoform are unable to transition from spermatogenesis to oogenesis. Finally, we performed similar experiments on *kah93* mutants, which lack the D isoform and have a truncated B isoform. These animals showed an intermediate phenotype, with normal number of sperm but reduced number of oocytes (Figure 5D and E). The reduced activity of the B isoform due to its truncation potentially allows other factors to transition the animals to oogenesis, resulting in the milder defects found in the *kah93* animals (**Figure 3B**).

**Figure 5.**
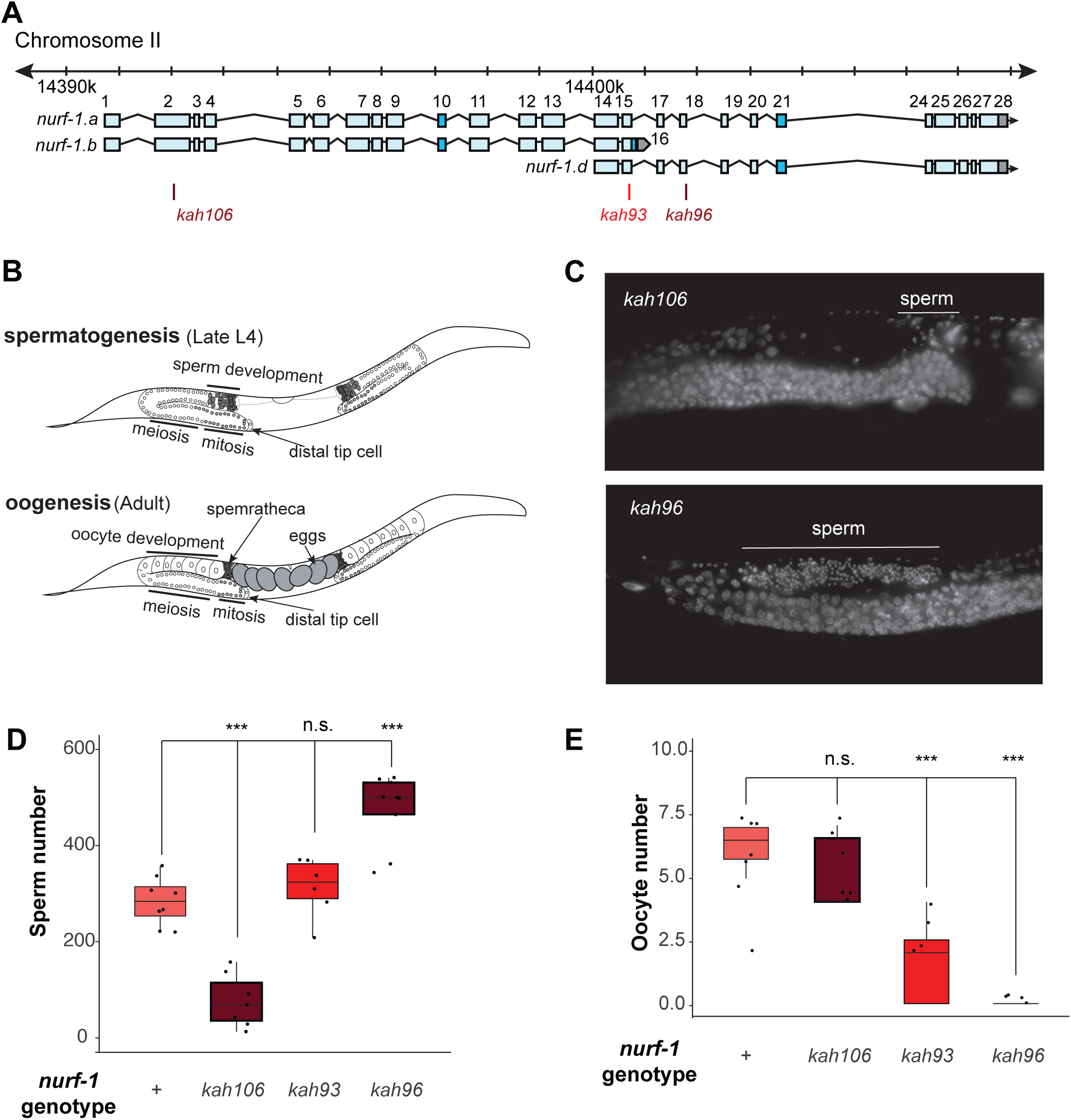
NURF-1.B and NURF-1.D have opposite effects on the sperm-to-oocyte switch in hermaphrodites. **A**) Genomic position of the previously-described stop codon mutants used in B and C. **B**) Summary of gametogenesis of *C. elegans*. Animals undergo spermatogensis during the late L4 and then transition to oogenesis stage during maturation to adulthood. The number of sperm produced during spermatogenesis can be determined by counting sperm in the spermatheca when oogenesis has begun. **C**) Represetative flourescence images of one spermatheca for DAPI stained young adult animals. Each tiny dot represents the condensed chromosomes of a single sperm. **D**) Sperm number of indicated strains. L4 animals were synchronized and allowed to develop for an additional 12 hours. DAPI staining was used to identify and count the number of sperm in each animal. Each dot represents a single animal. **E**) Oocyte number of indicated strains. L4 animals were synchronized and allowed to develop for an additional 12 hours. DAPI staining was used to identify and count the number of oocytes in each animal.

Although animals that lack either the B or D isoform are unable to reproduce, the cause of sterility is different at the cellular level. To further study the molecular effects of perturbing *nurf-1* function, we transcriptionally profiled adult N2*, NIL*_(nurf-1,LSJ2>N2*)_*, ARL_(del, LSJ2, N2*)_, and LSJ2 animals, which contain various combinations of the N2 and LSJ2-derived *nurf-1* mutations (**Table S1**). A multi-dimensional scaling plot indicated that the N2* and ARL_del_ replicates formed two unique clusters, and the LSJ2 and NIL*_nurf-1_* replicates largely overlapped in a third cluster (**Figure S12A**). The genetic variation surrounding the *nurf-1* locus is responsible for the majority of transcriptional differences between adult LSJ2 and N2* animals, suggesting most of the fixed variants do not have a dramatic effect on transcription on N2-like growth conditions. Although the LSJ2-derived 60 bp deletion regulates transcription, additional genetic variation in the NIL*_nurf-1_* strain, presumably from the N2-derived intron variant, also regulates transcription in adult animals.

To study the effects of the 60 bp deletion and intron SNV on transcription, we focused on two comparisons:

1. the N2* vs ARL_(del, LSJ2>N2*)_, which will identify transcriptional changes caused by the 60 bp deletion and
2. the NIL_(nurf-1, LSJ2>N2*)_ vs ARL_(del, LSJ2>N2*)_, which will identify transcriptional changes caused by the intron SNV (as well as linked mutations in the NIL <u>other than</u> the 60 bp deletion). We believe that the latter comparison will mostly report the changes of the intron SNV, as it accounts for most of the fitness differences between the two strains. We observed a large negative correlation between these two comparisons (**Figure S12B**). The most parsimonious explanation for this observation is that both the N2 and LSJ2-derived alleles in *nurf-1* regulate the activity of a common molecular target, which is likely to be the NURF complex.

### A duplication in a sister clade of *Caenorhabditis* species creates two separate *nurf-1* genes

Previous work in *C. briggsae* characterized the role of *nurf-1* in reproduction, including the isolation of *nurf-1* cDNAs in this species (Chen, Shen et al. 2014). Interestingly, although transcripts matching the *nurf-1.b* and *nurf-1.d* were isolated from this species, they no longer shared any exons with each other, suggesting that they were expressed from two separate genes (**Figure 6A**). We compared the gene products using BLAST and found that the shared exons in *C. elegans* had duplicated in *C. briggsae*, with one set of each retained in each of the new genes (**Figure 6A**). Short-read transcriptomics data for this species matched the cDNA analysis; we found evidence for transcripts matching *nurf-1.b, nurf-1.d,* and *nurf-1.f* (**Figure S13, S14**, and **S15**). Unlike *C. elegans*, *C.briggsae* seemed to have lost both the *nurf-1.a* and *nurf-1.q* transcripts.

**Figure 6.**
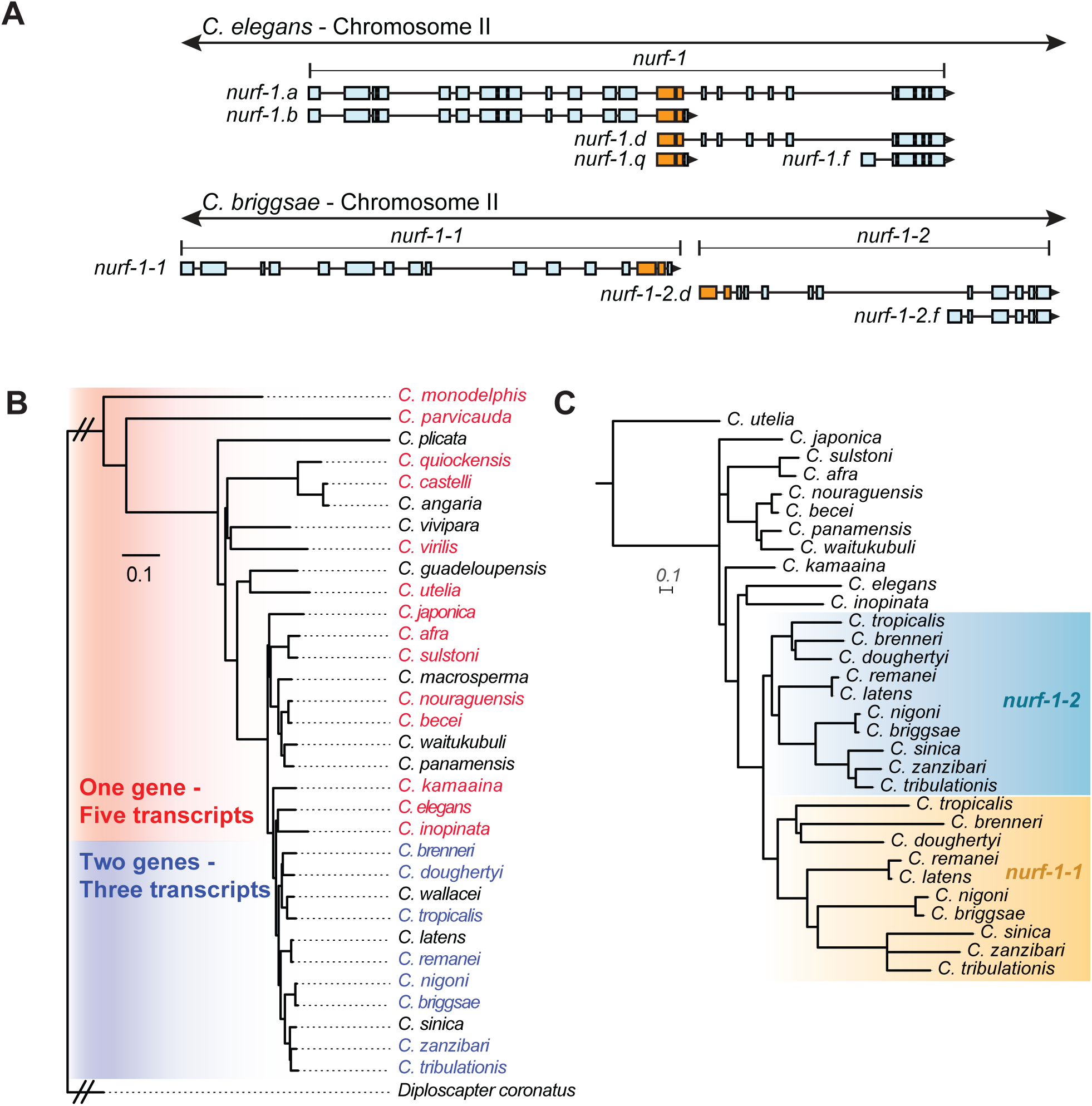
A duplication of the shared exons of the *nurf-1.b* and *nurf-1.d* transcripts resulted in the split of *nurf-1* it into two separate genes in a subclade of *Caenorhabditis* species. **A**) Comparison of two species with different versions of *nurf-1*. In *C. elegans*, *nurf-1.b* and *nurf-1.d* overlaps in the 14th and 15th exon (shown in orange). In *C.briggsae*, a duplication of the orange exons resulted in separation of *nurf-1.b* and *nurf-1.d* into separate genes*. C. briggsae* has also lost expression of the *nurf-1.a* and *nurf-1.q* transcripts. **B**) Distribution of the two versions of *nurf-1* (shown in panel **A**) in 32 *Caenorhabditis* species. Red indicates the *C. elegans* version, blue indicates the *C. briggsae* version, and black indicates a *nurf-1* version that could not be determined. The species phylogeny suggests that a duplication event occured in the common ancestor of the *brenneri/tribulationis* clade. **C)** The most well supported topology of the duplicated region is consistent with a single duplication event. Orange indicates protein sequence from the duplicated region in the *nurf-1-1* gene, and turquoise indicates protein sequence from the duplicated region in the *nurf-1-2* gene. Non-colored branches indicate unduplicated *nurf-1* sequence. The rate of amino acid substitution in the *nurf-1-1* duplicated region has also increased, as seen in the branch lengths. Scale is in substitutions per site.

Analysis of the *nurf-1* gene structure within the context of the *Caenorhabditis* phylogeny suggested that the exon duplication and separation of *nurf-1* into separate genes occurred at the base of a clade containing 11 described species, including *C. brenneri* and *C. tribulationis* (**Figure 6B**). We determined the *nurf-1* gene structure in 22 of the 32 *Caenorhabditis* species with published genomes and transcriptomes (Kiontke, Felix et al. 2011, Stevens, Félix et al. 2019) (**Figure S13, S14**, and **S15**). Like *C. briggsae*, the species in the *brenneri/tribulationis* clade express a transcript matching *nurf-1.b* from a single gene (which we call *nurf-1-1*). These species also express two transcripts matching *nurf-1.d* and *nurf-1.f* from a second gene, called *nurf-1-2*. None of these species appears to express *nurf-1.a* or *nurf-1.q* transcripts (Figure S13, **S14**, and **S15**). RNA-seq data for species outside of this clade (**Figure S13, S14**, and **S15**) matched the transcription pattern of *C. elegans,* suggesting that these species express five major transcripts from a single *nurf-1* gene: *nurf-1.a, nurf-1.b, nurf-1.d, nurf-1.f,* and *nurf-1.q.* These data suggest that the *C. elegans* transcript structure is ancestral.

Phylogenetic analysis of the duplicated ∼200 amino acid sequence was used to evaluate different hypotheses surrounding the timing and number of duplication events. The analysis supported the model that the split of *nurf-1* into two distinct genes happened once within the common ancestor of the *brenneri/tribulationis* clade (**Figure 6C –** additional possible trees shown in **Figure S16)**. The topology recovered for the region of *nurf-1* outside the duplication is congruent with the species tree (**Figure S17**). Interestingly, the rate of amino acid substitution in the duplicated region was accelerated in *nurf-1-1,* suggesting that this region experienced positive selection and/or relaxed selection after this duplication event occurred.

## Discussion

In this paper, we make use of CRISPR/Cas9-enabled gene editing to characterize the *nurf-1* gene in *C. elegans* and then study the sequence and expression of *nurf-1* orthologs in other *Caenorhabditis* species. The combination of genetics and evolutionary analysis allowed us to make a number of surprising observations. First, we show that an SNV in the 2^nd^ intron of *nurf-1* that fixed in the N2 laboratory strain increases the evolutionary fitness and fecundity of the N2 strain. Second, we show that the full-length isoform of *nurf-1* has split into two essential, mostly non-overlapping isoforms with opposite effects on cell fate in differentiating gametes. Finally, we show that the B and D isoforms have split into separate genes in a subset of *Caenorhabditis* species. These data show that *nurf-1* can be genetically perturbed to increase evolutionary fitness of animals in new environments and has experienced long-term evolutionary changes that have split its function and regulation into two isoforms/genes (Figure 7A and B).

**Figure 7.**
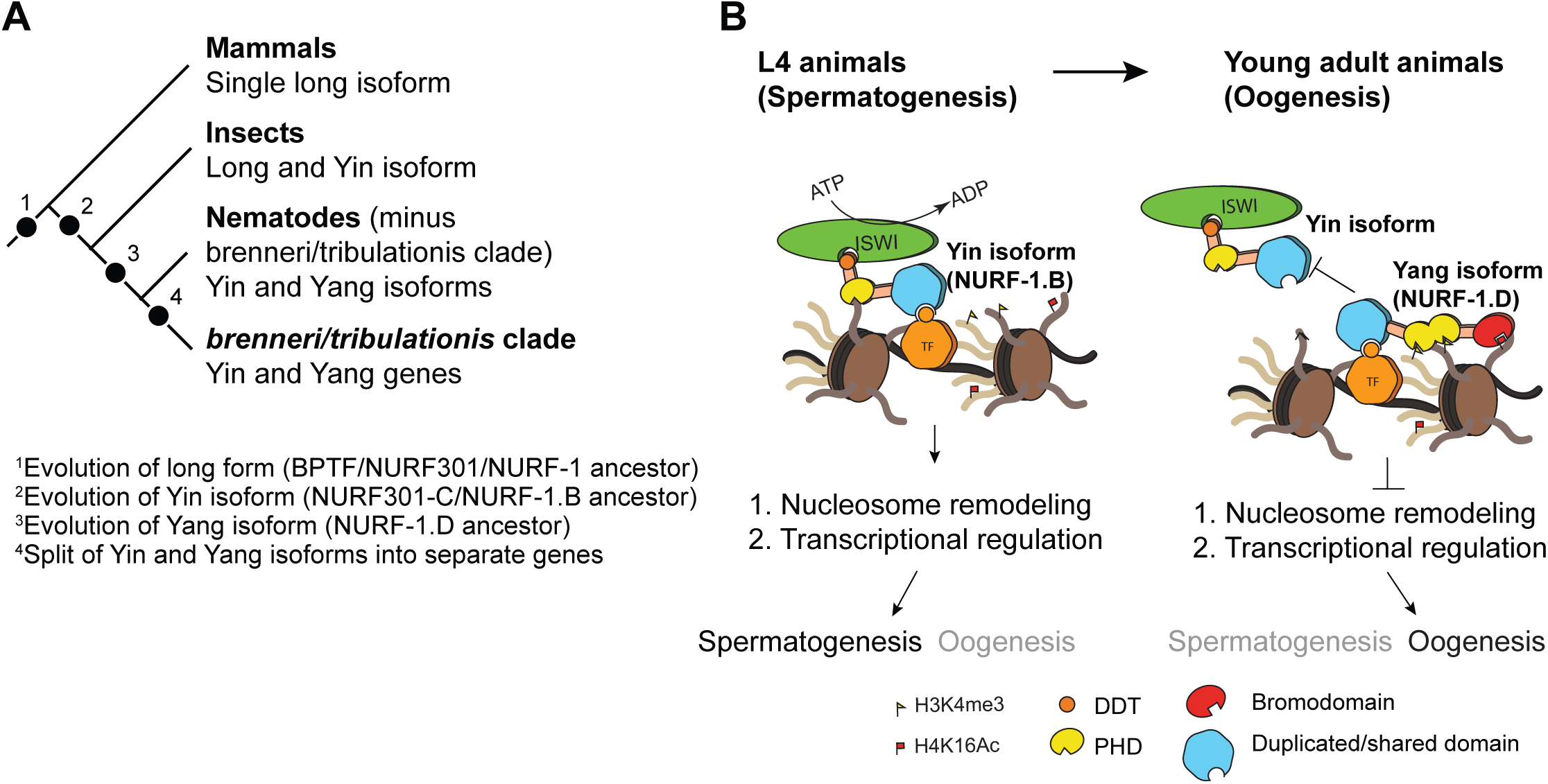
Proposed antagonistic (Yin-Yang) working model of two *nurf-1* isoforms in *C. elegans*. **A**) Descriptive phylogeny with proposed major transitions in *nurf-1* isoform evolution. Each dot indicate the timepoint of a major *nurf-1* isoform evolution event. **B**) Proposed molecular mechanism for NURF-1 isoforms. The NURF-1.B isoform interacts with ISWI through its DDT domain to form a NURF complex capable of remodeling chromatin at specific regions of the chromosome. NURF is recruited to these regions through interactions with specific transcription factors using protein domains encoded by the overlapping exons. This remodeling is necessary for transcriptional responses for spermatogenesis. Due to some unknown signal, after spermatogenesis has resulted in the production of ∼300 sperm, the NURF-1 D isoform outcompetes the NURF complex away from its target loci, casued the loss of transcription of key spermatogenesis genes, resulting in gametogenesis transitioning from spermatogenesis to oogenesis. The PHD and Bromodomain’s binding affinity to histone strengthens this repression, but they are not completely necessary for the ability of the D isoform to outcompete the B isoform.

### Evolution of NURF-1/BPTF across phyla

In humans and *Mus musculus*, an abundance of evidence confirms that the long-form isoform of *BPTF*, which is orthologous to *nurf-1*, is the primary isoform in the NURF chromatin remodeling complex (Alkhatib and Landry 2011). While a subset of *BPTF* exons are alternatively spliced, these events will not lead to the large changes in size we observe in the *nurf-1* gene. One exception is the FAC1 isoform, which encompasses 801 N-terminal amino acids of BPTF (Bowser, Giambrone et al. 1995). While FAC1 is found in amyloid Alzheimer’s patients and enriched in the nervous system (Bowser, Giambrone et al. 1995, Landry, Sharov et al. 2008), a biological role for this isoform has not been described. FAC1 is smaller and lacks conserved protein sequence found in the B isoform of *nurf-1,* suggesting an independent evolutionary origins and function.

In *Drosophila*, an intermediate state between humans and nematodes is found. Two major isoforms of NURF301 (the ortholog to *nurf-1*) have been described: a full-length form of NURF301 analogous to the full-length mammalian BPTF and an N-terminal form of NURF301 analogous to the NURF-1.B isoform of *C. elegans*. Both isoforms form NURF complexes and regulate gene expression (Kwon, Xiao et al. 2009). Genetic analysis suggests that full-length NURF301 is required for gametogenesis in both sexes while the N-terminal isoform is required for regulation of pupation and innate immunity.

Nematodes have retained the N-terminal isoform but seem to have lost use of the full-length isoform for most biological traits (Andersen, Lu et al. 2006). Instead, they express two C-terminal isoforms (D and F) that appear to be a recent evolutionary innovation, likely occurring before the origin of the *Caenorhabditis* lineage. We show that the D isoform (or the Yang isoform) is essential in *C. elegans*, and seems to act in opposition to the B isoform (or the Yin isoform) to regulate the sexual fate of differentiating gametes. The requirement of two antagonistic isoforms (the B and D) for reproduction is reminiscent of the principal of Yin and Yang. Genetic pathways often include both positive and negative regulators of transcription and ultimate phenotype, however, rarely are both the factors encoded by the same genetic locus. While there is growing appreciation of isoform-specific regulation of many genes, *nurf-1* appears to be unusually complex in this regard (although not unprecedented – see (Muller and Basler 2000, Berry, Miura et al. 2001, Wang, Xin et al. 2009).

We propose a molecular mechanism to explain the actions of the B and D isoforms to regulate transcription (**Figure 7B**). These two isoforms share 207 amino acids of protein sequence, which falls in a region that is thought to facilitate physical interactions with transcription factors (Alkhatib and Landry 2011). NURF-1.B participates as part of the NURF complex, which is recruited to certain promoters by binding to transcription factors. At these loci, NURF promotes or represses expression of target genes by remodeling the chromatin surrounding promoters and gene bodies. For unknown molecular reasons, NURF-1.D preferentially binds to these transcription factors, displacing the NURF complex from these genomic regions, causing a change of chromatin state and gene expression.

### Microevolution of NURF-1/BPTF

We showed that independent, beneficial alleles in *nurf-1* were fixed in two laboratory strains of *C. elegans* that each experienced an extreme shift in environment from their natural habitats. The N2-derived SNV results in the change of a run of homopolymers in the 2^nd^ intron of the *nurf-1.b* transcript. Such a change could act as an enhancer for the *nurf-1.d* promoter, but the nature of the genetic change and position is more consistent with a role in regulating the *nurf-1.b* transcript. Analysis of RNA sequencing data did not identify any obvious changes in levels of any of the *nurf-1* transcripts and it is unclear by what molecular mechanism it regulates *nurf-1* activity. Potentially, it could increase pausing of the RNA polymerase at the homopolymer run or could regulate RNA splicing by changing the secondary structure of the RNA molecule. In general, such a mutation would not be predicted by most bioinformatic approaches to have a phenotypic effect. Only the low genetic diversity between the LSJ2 and N2 strains allowed us to focus on this variant, and eventually demonstrate this particular variant is causal.

The probability of two beneficial mutations happening in both lineages by random chance is quite small. Less than 300 genes harbor derived mutations in either the N2 or LSJ2 lineage (McGrath, Xu et al. 2011). Only a handful of these fixed mutations are expected to be beneficial; our recent QTL mapping of fitness differences on agar plates only identified the *nurf-1* locus (Zhao, Wan et al. 2019) and the small effective population sizes (∼4-100) are expected to lead to the fixation of a number of nearly-neutral mutations through genetic drift and draft. Our work suggests *nurf-1* is a genetic target for adaptation to the extreme changes in environments associated with laboratory growth.

Targeting of *nurf-1* is consistent with its role as a regulator of life history tradeoffs. Many traits influence individual and offspring survival; however, the mapping of these traits onto fitness is thought to be dependent on the environmental niche an organism occupies. The LSJ2-derived deletion in *nurf-1* modified life history tradeoffs to prioritize individual survival over reproduction; by shunting energy away from reproduction and growth, they increased their chances of surviving on the poor, unnatural food. N2 animals grew on agar plates seeded with *E. coli* bacteria, which they can readily consume and metabolize into a useful energy source. In these conditions, survival is not the primary concern; each animal has three days to eat as much food as possible and produce as many progenies as possible to maximize the probability one of their offspring is transferred to the new food source. It is reasonable to think that the N2 and LSJ2 laboratory conditions represent opposite extremes along a life history axis encompassing individual survival and reproduction. The N2 mutation favors reproduction while the LSJ2 mutation favors survival.

In humans, alterations of *BPTF* are often observed in certain cancers. Tumor cells show a wide range of phenotypes, including proliferation and quiescence. Genetic alterations of BPTF in cancer evolution might shift cells between these life history strategies.

### Split of *nurf-1* into separate genes potentially resolves conflict between the Yin and Yang isoforms caused by shared exons

Our work suggests that genetic changes in *nurf-1* can be selected by short-term adaptation. As organisms evolve, recruitment of NURF to specific loci could be accomplished by changing its binding with specific transcription factors through amino acid changes in NURF-1. The most rapidly evolving portion of the protein is within the 14^th^ and 15^th^ exons, suggesting positive selection acts on this region of the protein, potentially changing the transcription factors NURF-1 binds to. One potential issue that arises in this situation is the pleiotropy of genetic changes in the shared region; changing the amino acid sequence of the B isoform also changes the D isoform. Are there situations where modifying one isoform but not the other is preferred? Escape from adaptive conflict is a mechanism by which gene duplication can resolve the situation where a single gene is selected to perform multiple roles (Des Marais and Rausher 2008). After duplication, each copy is free to improve its function independently.

In a clade of *Caenorhabditis* nematodes, the *nurf-1* gene has split into two separate genes in a manner consistent with escape from adaptive conflict. Duplication of the shared exons releases each isoform to evolve independently. Our data suggests that after this event, the duplicated region in the *nurf-1-1* gene experienced accelerated evolution, consistent with an increased rate of adaptive evolution.

This duplication event also resolved a molecular conflict between the *nurf-1.b* and *nurf-1.d* transcripts. To produce both transcripts in the same cell, there must be a mechanism to distinguish between transcripts containing the 1^st^ to 15^th^ exons (the *nurf-1.b* transcripts) and transcripts initiating from the 14^th^ exon (the *nurf-1.d* transcript). In the former case, the 15^th^ exon is spliced to the 16^th^ exon to terminate the transcript. In the latter case, the 15^th^ exon is spliced to the 17^th^ exon and continues transcription. Alternatively, the cell might not distinguish between transcripts, but uses each alternative splice site at a constant ratio (i.e. 80% of the time, the 15^th^ exon is spliced to the 16^th^ exon and 20% of the time, the 15^th^ exon is spliced to the 17^th^ exon). In the latter scenario, two additional transcripts must be produced. Intriguingly, these two transcripts match *nurf-1.a* and *nurf-1.q,* suggesting these transcripts are non-functional biproducts of molecular conflict between *nurf-1.b* and *nurf-1.d*.

Multiple lines of evidence are consistent with the 2^nd^ scenario. First, while the *nurf-1.q* transcript is produced at high levels, we were unable to observe its product in our immunoblots, suggesting that it is either not translated or the protein product is rapidly degraded. Second, our genetic tests were unable to identify a biological role for *nurf-1.a.* Third, we observe a loss of both the *nurf-1.a* and *nurf-1.q* transcripts in the species that have split *nurf-1* into two genes. It would have been quite easy for these species to retain expression of *nurf-1.q* in their current configuration, either through a promoter in front of the 14^th^ exon in the *nurf-1-1* gene, or an alternative stop exon after the 2^nd^ exon of the *nurf-1-2* gene, since both of these elements existed in the ancestral state.

If this scenario is true, there are a few interesting implications. The most highly expressed *nurf-1* transcript, *nurf-1.q*, might simply be a non-functional biproduct of a molecular conflict in splicing. Transcript levels are often used as a proxy for biological function, however, our work suggests that high expression of a transcript could also occur as a mechanistic bioproduct of splicing. This model also suggests a molecular signature useful for identifying other genes in similar conflict – the production of four transcripts from two promoters sharing an alternative splice site. As long read technology matures, accurate characterization of transcripts will enhance our ability to understand isoform production in a large number of species.

## Conclusion

A fundamental problem in evolutionary biology is understanding the genetic mechanisms responsible for phenotypic diversity in extant species. Here, we present one route to address this problem. Experimental evolution and genetic analysis can be used to identify evolutionary relevant genes and understand their function. This knowledge can be leveraged to understand patterns of evolution of these genes in other species. We believe that merging genetics, genomics, and molecular evolution is a powerful approach to understand the evolutionary mechanisms responsible for long-term adaptation and species level differences.

| Name | Evidence |  | Size |  | Conserved<sup>c</sup> | Predicted biological role in *C. elegans*<sup>d</sup> | Other names |
| --- | --- | --- | --- | --- | --- | --- | --- |
|  | Transcript<sup>a</sup> | Protein<sup>b</sup> | aa | kD |  |  |  |
| *nurf-1.a* | N | - | 2197 | 252 | M,D | None | Full-length |
| *nurf-1.b* | C,N,I | W | 1621 | 186 | D | Reproduction, vulval development | N-terminal or NURF-1.A |
| *nurf-1.d* | C,N,I | W | 816 | 92 | - | Size, dauer, reproduction, axon guidance | C-terminal or NURF-1.C |
| *nurf-1.f* | C,N,I | W | 581 | 58 | - | None | NURF-1.E |
| *nurf-1.q* | N,I | - | 243 | 36 | - | None | - |

<sup>a</sup> C indicates full-length cDNA have been isolated for this transcript, N indicates evidence from direct sequencing of RNA or cDNA using Oxford Nanopore reads support this transcript, and I indicates evidence from Illumina short read RNAseq supports this transcript
<sup>b</sup> W indicates evidence for the protein isoform was obtained using Western blot
<sup>c</sup> M or D indicate an analogous isoform is described in mammals (mice or humans) or *Drosophila*, respectively
<sup>d</sup> Predictions from Andersen et al (2006), Large et al (2015), or Mariana et al (2016)

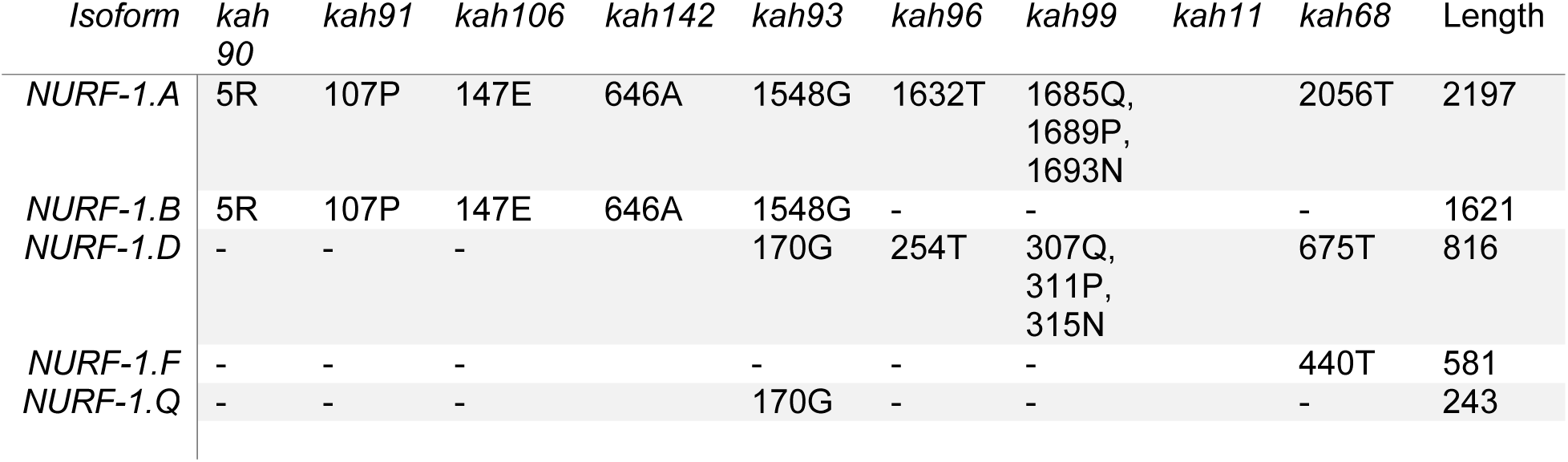

## Acknowledgements

We thank the *Caenorhabditis* Genetics Center, which is funded by NIH Office of Research Infrastructure Programs (P40 OD010440), for strains, and WormBase for information. We are grateful to Rachael Workman and Winston Timp for sharing Oxford Nanopore reads of *nurf-1* prior to publication. We thank Matthew Rockman, Luke Noble, Janna Fierst, Erich Schwarz, and Janet Young for access to unpublished genomic data. We also thank Todd Streelman, Greg Gibson, Soojin Yi, David Katz, Annalise Paaby, and members of the McGrath lab for discussions, and Annalise Paaby and Erik Andersen for comments on the manuscript. This work was supported by NIH R01GM114170 (to P.T.M.) and R01GM121688 (to R. E. E.).

## Methods

### Strains

The following strains were used in this study:

#### Near isogenic lines (NILs)

CX12311 (N2*): *kyIR1(V, CB4856>N2), qgIR1(X, CB4856>N2)*,

PTM66 (NIL_(*nurf-1*,LSJ2>N2*)_): *kyIR87(II, LSJ2>N2); kyIR1(V, CB4856>N2), qgIR1(X, CB4856>N2)*

#### CRISPR-generated allelic replacement lines (ARLs)

PTM88 (ARL_del, LSJ2>N2_): *kyIR1(V, CB4856>N2); qgIR1(X, CB4856>N2); nurf-1(kah3)II; spe-9(kah132)I*

PTM416 (ARL_intron,LSJ2>N2_): *nurf-1(kah127)II*

PTM417: *kyIR1(V, CB4856>N2); qgIR1(X, CB4856>N2); nurf-1(kah3)II*

#### CRISPR-generated barcoded strains

PTM229: *dpy-10(kah82)II*

PTM288: *kyIR1(V, CB4856>N2); qgIR1(X, CB4856>N2); dpy-10(kah82)II*

#### CRISPR-generated epitope-tagged strain

PTM420 (HA-FLAG): *nurf-1(kah124,kah133)II,*

#### CRISPR-generated STOP codons replacement lines

PTM98 (exon23): *nurf-1(kah11)II*

PTM203 (exon26): *nurf-1(kah68)II*

PTM316 (exon 1): *nurf-1(kah90)II/ oxTi924 II*

PTM317 (exon 2): *nurf-1(kah91)II/ oxTi924 II*

PTM319 (exon 15): *nurf-1(kah93)II/ oxTi924 II*

PTM322 (exon 18): *nurf-1(kah96)II/ oxTi924 II*

PTM325 (exon 19): *nurf-1(kah99)II/ oxTi924 II*

PTM332 (exon 2): *nurf-1(kah106) II/ oxTi924 II*

PTM487 (exon 7): *nurf-1(kah142) II/oxTi721 II*

#### CRISPR-generated domain replacement lines

PTM113 (PHD1): *nurf-1(kah16)II*,

PTM116 (PHD2): *nurf-1(kah19)II*,

PTM117 (PHD2): *nurf-1(kah20)II*,

PTM118 (Bromodomain): *nurf-1(kah21)II*,

PTM167 (Bromodomain): *nurf-1(kah32)II*,

PTM170 (double PHD): *nurf-1(kah19,kah36)II*,

PTM189 (3 domains): *nurf-1(kah19,kah36,kah54)II*,

PTM211 (double PHD): *nurf-1(kah66,kah73)II*

#### MosSCI transgenic strains

PTM371: *nurf-1(kah93) II/ oxTi721 II; kahSi7*,

PTM372: *nurf-1(kah96) II/ oxTi721 II; kahSi7*,

PTM373: *nurf-1(kah99) II/ oxTi721 II; kahSi7*,

PTM376: *nurf-1(n4295) II; kahSi7*,

PTM517: *kyIR1 (V, CB4856>N2); qgIR1 (X, CB4856>N2); nurf-1(kah3) II; kahSi7*

#### CRISPR-generated deletion strains

PTM512 (23^rd^ exon deletion): *nurf-1(kah149) II*

PTM489 (HA-FLAG + 23^rd^ exon deletion): *nurf-1(kah124,kah133,kah144)II*

#### Other double mutants

PTM354: *nurf-1(n4295, kah113) II/ oxTi924 II*

### Strain construction

#### Previously described strains

CX12311, PTM66, and PTM88 were all previously described (McGrath, Xu et al. 2011, Large, Xu et al. 2016).

#### CRISPR-generated allelic replacement lines (ARLs)

We used the coCRISPR protocol to generate all CRISPR-edited lines using single-strand oligonucleotides to make precise edits (Arribere, Bell et al. 2014, Paix, Folkmann et al. 2015).

Resequencing of the PTM88 strain identified a number of background mutations, including an A to G missense SNV that is predicted to change an asparagine to an aspartic acid which we named *kah132*. The flanking sequence of this mutation is 5’-cgacaatgac[a]atcgccaggg-3’. We backcrossed out this *spe-9(kah132)* mutation, along with additional background mutations, to create PTM417.

To create PTM416, we designed a number of guide RNAs nearby the intron SNV. However, we were unable to identify editing events using these guide RNAs, putatively due to the high usage of As and Ts.

We turned to a two-step strategy to create the edit, first creating a deletion of the 2^nd^ intron along with flanking exon regions using guide RNAs with high predicted efficiency. We created the following constructs driving the following sgRNAs:

5’- TCGATAATTATCCGTTTGT(GGG) -3’,

5’- TTGCATCATATCCCACAAA(CGG) -3’,

5’- ACGGTAGCTCATGAAGAGA(AGG) -3’

and 5’- TTCCGACGAATATAAGAAA(CGG) -3’

We also ordered an oligonucleotide repair:

5’- GTCTGTTAGAGATGCTATTAATGTCGATAATTATCGCTACCATAGGCACCACGAGCGAGATTCGTCGGAATTTAAGAAACTTGTGAATAATGTT -3’

We injected 50 ng/μl of P*_eft-3_*::Cas9, 25 ng/μl of *dpy-10* sgRNA, 500 nM *dpy-10(cn64)* repair oligo, 10 ng/μl of each of the *nurf-1* sgRNAs listed above, and 500 nM of the repair oligonucleotide into CX12311 animals.

Jackpot broods were identified and roller animals were genotyped using the following primers along with the BanI restriction enzyme:

5’- GCAGGCCGGCCTTCGCGCCTGGGTAATACC -3’ and

5’- CGGCAGTTTTCGTCGTTCTG -3’

A single heterozygote worm was identified. Wild-type heterozygote progeny were identified (to remove the linked *dpy-10* mutation) and this mutation was balanced (homozygous animals were sterile) with an integrated GFP marker near the *nurf-1* gene (*oxTi924*). This strains was frozen with the following genotype: PTM366 *nurf-1(kah125)/oxTi924* II; *kyIR1 (V, CB4856>N2); qgIR1 (X, CB4856>N2) X*.

For the second step, we crossed PTM366 to PTM66 animals and selected non-fluorescing animals to create *nurf-1(kah125)/kyIR87(II, LSJ2>N2)*; *kyIR1 (V, CB4856>N2); qgIR1 (X, CB4856>N2) X* compound heterozygote animals. We used the following sgRNAs to specifically target the nurf-1(kah125) homologous chromosome:

5’- ATCTCGCTCGTGGTGCCTA(TGG) -3’

and 5’- TTCCGACGAATCTCGCTCG(TGG) -3’

The 2^nd^ homologous chromosome, containing the *kyIR87* introgression was used as a repair construct. We injected 50 ng/μl P*_elt-3_*::Cas9, 10 ng/μl *dpy-10* sgRNA, 500 nM *dpy-10(cn64)* repair oligo and 25 ng/μl of each *nurf-1* sgRNA. Roller animals were then PCR genotyped to screen for animals that were homozygous for the LSJ2 allele at the intron and heterozygote for the 60bp deletion.

After screening, the target genotype was made homozygote. This strain was named PTM410 *kyIR1 (V, CB4856>N2); qgIR1 (X, CB4856>N2); nurf-1(kah127)II*. PTM416 was created by backcrossing the PTM410 strain to the N2 background using an RFP fluorescent *nurf-1* balancer (*oxTi721)* strain for 4 generations. We genotyped the *npr-1* and *glb-5* sites to verify that PTM416 did not carry the introgressions surrounding these genes.

#### CRISPR-generated isotope-tagged lines

To create the PTM420 epitope-tagged strain the following guide RNA and repair oligo was used to first add an HA epitope tag into the 16^th^ exon:

5’-TGGCACTTGCTCAGTTGTGG-3’

5’-TTTTGTCAAATTTGGAGCCGTTTGGGGAACCTCTAGGCGTAGTCGGGGACGTCGTATGGGTATCCTCCTCCTCCTCCTCCCTGCTGTTCGTCTGGGACCTGCTCGGTTGTAGTAGAAACTGCGAAACCAGTCGCGTCATCAGGCATGTC-3’

The following injection mix was used: 50 ng/μl P*_eft-3_*::Cas9, 10 ng/μl *dpy-10* sgRNA, 500 nM *dpy-10(cn64)* repair oligo, 25 ng/μl of sgRNA, and 500 nM repair oligonucleotide.

We next added a 3xFLAG tag to the C-terminal of *nurf-1* gene using purified Cas9 protein (IDT, Catalog #1074181) and *in vitro* synthesized RNAs (Synthego) using a modified protocol (Prior, Jawad et al. 2017). The injection mix was prepared as follows: 2uM *dpy-10* sgRNA (RNA scaffold 5’- GCUACCAUAGGCACCACGAG -3’ + tracrRNA) and 4uM of two sgRNAs that targeted this region (RNA scaffold: 5’- CUCAUAAGUUCGCAUCCAG -3’+ tracrRNA, 5’- UUCGGAUCAGCUGUUGCCAC -3’+ tracrRNA) were mixed and incubated in a thermocycler at 95°C for five minutes, then 2.5ug/ul Cas9 protein was added and incubated at room temperature for five minutes. Finally, 0.2uM *dpy-10* repair oligo and 0.5uM FLAG repair oligo were added to mix and incubate at room temperature for 60 minutes. This mix was injected into the HA-tagged strain to create the double epitope tagged line.

#### CRISPR-generated STOP codon replacement lines, PHD/bromodomain replacement lines, and deletion

The following injection mix was used to create each of these strains: 50 ng/μl P*_eft-3_*::Cas9, 10 ng/μl *dpy-10* sgRNA, 500 nM *dpy-10(cn64)* repair oligo, 25 ng/μl of sgRNA, and 500 nM repair oligonucleotide. For each strain/allele, each of the specific sgRNAs and repair oligos used to construct it are listed in **Table S4**. To facilitate the genotyping process, some of the repair oligos for STOP codon replacement sites contain restriction sites that will alter some of the amino acids, exact changes are listed in **Table S4**. In *C. elegans* nomenclature, Identical edits must be given different allele names if they were isolated independently.

For mutants that were sterile (or lead to sterility), we balanced these mutations using a GFP (*oxTi924*) or mCherry (*oxTi721*) integrated marker near *nurf-1*.

#### MosSCI transgenic strains

MosSCI strain construction was done following standard protocol from Frøkjær-Jensen et. Al (Frokjaer-Jensen 2015). Injection mix was prepared as following: 38ng/ul pCFJ601 (Mos1 transposase), 30ng/ul pCFJ151 - P*nurf-1.d::nurf-1.d-SL2-GFP* (insertion vector with homologous arms), 2.5ng/ul pCFJ90 (*Pmyo-2*::mCherry), 5ng/ul pCFJ104). This was injected into EG6699 uncoordinated animals. Three injected animals were placed on a single plate at 30 °C to facilitate starvation. After 5 days, coordinated animals with GFP fluorescence and no red fluorescence were singled to new NGM plates and allowed to proliferate. Their progenies were singled and a single homozygote without uncoordinated offspring was maintained. This homozygote was then backcrossed to N2 for 4 generations to remove *unc-119(ed3) III* to create the PTM337 strain containing the integrated rescue construct. This strain was then crossed to a variety of *nurf-1* alleles using standard protocols.

### Molecular biology

All sgRNAs were constructed using NEB Q5 site directed mutagenesis kit (E0554) using primers 5’- [unique sgRNA protospacer sequence] + GTTTTAGAGCTAGAAATAGCAAGT -3’ and

5’- CAAGACATCTCGCAATAGG -3’

to modify a vector backbone containing a subclone of pDD163 containing the U6 promoter to drive sgRNAs in germline^1^.

To create the pCFJ151 - P*nurf-1.d::nurf-1.d-sl2-GFP* plasmid, a *nurf-1.d* cDNA was isolated from reverse transcribed RNA using primers containing NheI restriction sites. This PCR product was then digested and ligated to a pSM vector. A 2890bp long promoter region immediately upstream of the *nurf-1.d* isoform was amplified with a forward primer including FseI and a reverse primer including AscI restriction sites. This PCR product was then digested and ligated into the vector constructed in step 1. Thrid, an SL2-GFP sequence from was cut and ligated into the new vector using KpnI and SpeI restriction sites. Finally, this entire sequence containing the promoter, cDNA and sl2::GFP sequence was inserted into the pCFJ151 vector using NEB Q5 site directed mutagenesis kit.

### Nematode growth conditions

The animals were cultured on 6cm standard nematode growth medium (NGM) plates containing 2% agar seeded with 200 μl of an overnight culture of the *E. coli* strain OP50. Growth temperature was controlled using a 20°C incubator. Strains were grown for at least three generations without starvation before any experiments was conducte.

### *nurf-1* conserved regions

The predicted protein sequence for the NURF-1.A protein isoform was blasted against human or *Drosophila melanogaster* protein databases using NCBI blastp (McGinnis and Madden 2004). Regions with alignment scores above 50 were annotated as homologous regions. These homologous regions were further verified through multiple sequence alignment in Clustal Omega program (Chojnacki, Cowley et al. 2017).

### Competition experiment

Competition experiments were performed as described previously (Zhao, Long et al. 2018).

### RNAseq analysis

#### RNAseq samples for comparing the effect of the nurf-1 intron SNV

N2 and PTM416 worms were synchronized using a 3-hour hatch-off. Worms were observed every hour after 46 hours until the majority were in the L4 stage (which occurred at 48 hours). Four hours later, worms were collected and kept frozen in −80 °C freezer until RNA extraction for the 52 hour timepoint. Eight hours later, young adult animals were collected and kept frozen in the −80 °C freezer until RNA extraction for the 60 hour timepoint.

#### RNAseq samples for comparing effect of the two derived nurf-1 mutations

CX12311, PTM66, PTM88, LSJ2 L4 hermaphrodites were picked to fresh NGM agar plates. Their adult progeny were bleached using alkaline-bleach solution to isolate eggs for synchronization. The eggs were washed with M9 buffer for three times and placed on a tube roller overnight. About 400 hatched L1 animals were placed on NGM agar plates and incubated at 20°C until they reach young adulthood, as determined by when eggs were observed on assay plates. These worms were then harvested, washed 3 times with M9 buffer, and frozen in a −80 °C freezer for later processing.

#### RNAseq samples for heat shock

N2 and PTM416 worms were synchronized using a 3-hour hatch-off. Eggs were cultured at 20°C until they reached L4 stage. Heat shock assay plates were then wrapped with parafilm and placed in a water bath pre-heated to 34°C for 2 hours or 4 hours. Worms were either collected right after heat shock or after 30 minutes at 20°C for the recovery group.

For each of the above experiments, RNA was isolated using Trizol. The RNA libraries were prepared using an NEB Next Ultra II Directional RNA Library Prep Kit (E7760S) following its standard protocol. The libraries were sequenced by an Illumina NextSeq 500. The reads were aligned by HISAT2 using default parameters for pair-end sequencing (Kim, Langmead et al. 2015). Theses aligned reads were then visualized in IGV browser (Robinson, Thorvaldsdottir et al. 2011) to examine *nurf-1* splice junction track (as shown in **Figure S3**). Transcript abundance was calculated using featureCount and then used as inputs for the SARTools. SARTools use edgeR for normalization and gene-level differential analysis (Varet, Brillet-Gueguen et al. 2016) and output the multidimensional scaling plot for each transcriptome analysis project. Differentially expressed genes were determined for comparisons have adjusted p-value < 0.05. Genes upregulated and downregulated are plotted separately for the tissue and stage analysis. Each gene is normalized by dividing the sum of its expression level across all stages. And this normalized table is used for hierarchical clustering analysis. Sequencing reads were uploaded to the SRA under PRJNA526473.

Kallisto was used to quantify abundances of *nurf-1* transcripts (Bray, Pimentel et al. 2016). We first created our own reference transcriptome by modifying the transcripts in Wormbase published reference transcriptome to restrict our analysis to the *nurf-1.a, nurf-1.b, nurf-1.d, nurf-1.f* and *nurf-1.q* isoforms. Alternative splicing sites in the 10^th^, 16^th^, and 21^st^ exons were also removed from this reference database to ensure they were consistent between all isoforms. We used wildtype L2 RNAseq data from Brunquell et. al to quantify wildtype *nurf-1* abundance (Brunquell, Morris et al. 2016) and extracted tpm(transcripts per million) data from Kallisto output abundance table. We used RNAseq data from PRJNA311958 and PRJNA321853 (Brunquell, Morris et al. 2016) (Li, Chauve et al. 2016) to quantify the heat shock response of *nurf-1* isoforms in **Figure S11B**.

### Western blot

4 N2 and PTM420 gravid hermaphrodites were picked to fresh 5.5cm NGM agar plates. Worms were collected just prior to starvation using M9 buffer and stored at −80°C until protein extraction. At least 4 plates of worms were used for each protein isolation. Worms were condensed by centrifugation and 2x sample buffer (100 mM Tris-HCl pH 6.8M, 200mM dithiothreitol, 4 % SDS, 0.2 % Bromophenol Blue, 20% glycerol) was added in 1:1 w/v ratio. 1ul of 500mM EDTA and 1ul of Halt protease inhibitor cocktail (100x) (Catalog number: 78430) were added for every 100ng of worm sample. The protein sample was vortexed for 90 seconds and incubated on ice for about 1 minute. Samples were then sonicated in a Bransonic 0.5 gallon ultrasonic bath filled with hot water > 80°C for 10 minutes and immediately placed on ice for 2 minutes. We then boiled the samples for 5 minute and placed on ice for cooling down. The sample was centrifuged at 12,000 rpm for 5 minutes and the supernatant was transfered to new tubes.

All samples were loaded on 5% SDS-PAGE gel at 3ul, 5ul and 7ul volumes followed by Coomassie blue staining and washing steps. Gels were then dried using DryEase Mini-Gel Drying System (Invitrogen, Catalog number: NI2387). These gels were used to normalize protein loading volume for different samples.

Each sample was loaded onto a freshly made 6% or 10% SDS-PAGE gel and run at 25mA. Gel samples were then transferred in 10mM CAPS pH10.5 buffer at 20V and 20mA for 17hrs to a PVDF membrane. Protein products with HA tag were detected using 1:500 anti-HA antibody (Life Technologies, Catalog number: 326700), NURF-1.D isoform with FLAG tag was detected using 1:1000 PIERCE ANTI-DYKDDDDK antibody (Life Technologies, Catalog number: MA191878) and NURF-1.F isoform with FLAG tag was detected using 1:1000 Millipore ANTI-FLAG antibody (Millipore Sigma, Catalog number: F3165).

### Egg-laying analysis

Egg laying assays were performed as previously described (Large, Xu et al. 2016). All egg-laying assays were carried out at 20°C using standard 3cm NGM plates seeded with the OP50 strain of Escherichia coli. OP50 were prepared freshly by streaking a glycerol stock of OP50 on an LB plate and letting grow at 37°C overnight. A single colony was then picked to 5ml fresh LB and cultured overnight in a shaking incubator at 200rpm. 1ml of the overnight culture was used to inoculate 200 ml of LB for 4–6 hours of growth at 37°C with shaking. The 200ml OP50 culture was concentrated via centrifugation to an OD600 of 2.0 and this culture was used for seeding experimental plates with 50 μl aliquots. All experimental plates were prepared the week of the assay and left at 22.5°C 18–24 hrs following seeding. Plates were then placed at 4°C until the day of the assay and warmed to 20°C for 12 hours before each time point.

For strains that have severe reduced fertility when homozygote, one L4 nematode was transferred to the 50μl experimental plate. The number of eggs laid were measured every 12 or 24 hours, and eggs laid per hour was calculated by dividing the time range and number of animals left on each plate at each timepoint. At least 10 replicates were assayed for each strain.

For other strains, six fourth larval stage (L4) nematode was transferred to the 50μl experimental plate. The number of eggs laid were measured every 12 or 24 hours, and eggs laid per hour was calculated by dividing the time range and number of animals left on each plate at each timepoint. Six replicates were assayed for each strain.

Fecundity was calculated by summing up all eggs laid for each worm.

### Growth analysis

For trains with mutation in PHD or Bromodomains, growth analysis were performed as previously described (Large, Xu et al. 2016). For other strains, video tracking was done the same way but data analysis was performed by painting each moving animals and measure the average area for each individual worms during the course of video tracking. For strains that were balanced with fluorescent markers, only non-fluorescent worms were picked for video tracking.

### Sperm and oocyte counting analysis

4 N2, PTM332, PTM319 and PTM332 gravid hermaphrodites were picked to fresh 5.5cm NGM agar plates. After 3 days, 20-30 non-fluorescent L4 worms were picked to a new NGM plate and let grow at 20°C for 12 hours. Worms were then picked to a drop of M9 buffer on a Fisher Superfrost Plus slide (22-037-246). Fixation was done through applying 95% ethanol for three times. A drop of Vector Laboratories Vectashield Mounting Medium with DAPI (H-1500) was added and a coverslip was applied and sealed with nail polish. Z-stack images were captured through a moving-stage Olympus IX73 microscope under 40x objective. Oocytes were counted while imaging and sperm number was measured manually by analyzing z-stack images on ImageJ through the CellCounter plugin.

### Genomic and Transcriptomic analysis of *nurf-1* in additional *Caenorhabditis* species

To identify *nurf-1* orthologs, we used homology information included in www.wormbase.org or by blasting *C. elegans* protein sequences against protein data provided by the *Caenorhabditis* genome project (http://blast.caenorhabditis.org). Genomic regions that contain the identified *nurf-1* orthologs and related gff3 annotation data were downloaded from <u>download.caenorhabditis.org</u>or the WormBase public FTP site (data from (Stein, Bao et al. 2003, Mortazavi, Schwarz et al. 2010, Fierst, Willis et al. 2015, Slos, Sudhaus et al. 2017, Kanzaki, Tsai et al. 2018, Yin, Schwarz et al. 2018, Lamelza, Young et al. 2019)). Species with public RNAseq data were identified in the SRA database. These reads were downloaded and aligned to corresponding *nurf-1* DNA reference sequence for each species using HISAT2 and further manipulated using SAMTOOLS (Li, Handsaker et al. 2009, Kim, Langmead et al. 2015). Gene annotations were manually corrected by inspecting the RNAseq predicted intron sequences and used to generate Sashimi plots using the IGV browser (Robinson, Thorvaldsdottir et al. 2011, Katz, Wang et al. 2015). The Sashimi Plot parameter Junction Coverage Min was adjusted for each species to best visualize the exon-exon junctions based upon coverage data. To identify the duplicated region for the NURF-1.B and NURF-1.D isoforms, we blasted each B isoform against a database of the D isoforms, and vice-versa. The homologous regions for each protein were refined using a multiple sequence alignment of NURF-1.B and NURF-1.D proteins using Jalview (Waterhouse, Procter et al. 2009). For some of the species that we were unable to resolve the full *nurf-1* region (due to missing sequence for part of the region), we were able to identify the duplicated region and included this in the phylogenetic analysis.

### Phylogenetic analysis

We aligned the protein sequences of the duplicated region from the *nurf-1* loci of 21 *Caenorhabditis* species using MAFFT (Katoh and Standley 2013). We also aligned the protein sequences for regions outside the duplicated region. Maximum likelihood trees were estimated for each alignment along with 1000 ultrafast bootstraps (Hoang, Chernomor et al. 2018) using IQ-TREE (Nguyen, Schmidt et al. 2015), allowing the best-fitting substitution model to be automatically selected (Kalyaanamoorthy, Minh et al. 2017). We noted that the resulting topology recovered for the duplicated region was incongruent with the species tree, likely due to limited phylogenetic signal in the short alignment (**Figure S18**). To address this, we instead assessed the levels of support for alternative phylogenetic hypothesis surrounding the number and timing of duplication events that we congruent with the species tree. Log-likelihoods were calculated for each topology and an approximately unbiased (AU) test (Shimodaira 2002) was performed using IQ-TREE. Newick trees were visualized using the iTOL web server (Letunic and Bork 2016).

### Statistics

Significant differences between two means were determined using two-tailed unpaired t-test. To correct for multiple comparison, we used Tukey multiple comparison test.

---

**Figure S1.**
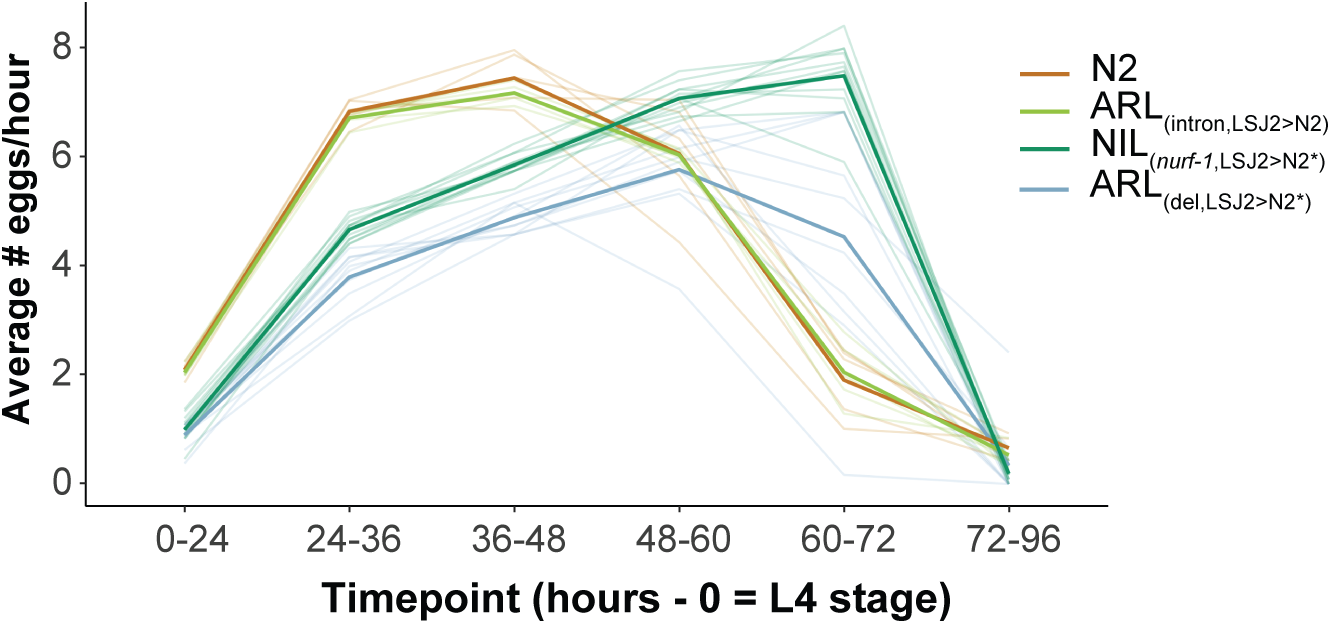
Egg-laying rate of four strains. Egg-laying rate was calculated at the indicated time points. Six L4 animals were picked onto each assay plates (time = 0) for the indicated times. The total number of eggs was counted for each plate and used to calculate the average number of eggs laid per hour for each animal. Each individual trial is shown as a dim line. The mean for each strain is shown as the bold, colored lines. But the effect of the intron SNV on this trait was subtle, especially when compared to the difference in reproductive rate caused by the *nurf-1* 60bp deletion (Figure S1). We believe this is explained by epistasis between the two *nurf-1* mutations, with the LSJ2 combination of both alleles having a non-linear effect on reproductive rate. We are unable to test this hypothesis as we did not construct a double ARL strain containing LSJ2 alleles of both of the *nurf-1* mutations due to the difficulty in creating the intron edit (see Methods). Alternatively, additional mutations in NIL_(*nurf-1*,LSJ2>N2*)_ could contribute to reproductive rate.

**Figure S2.**
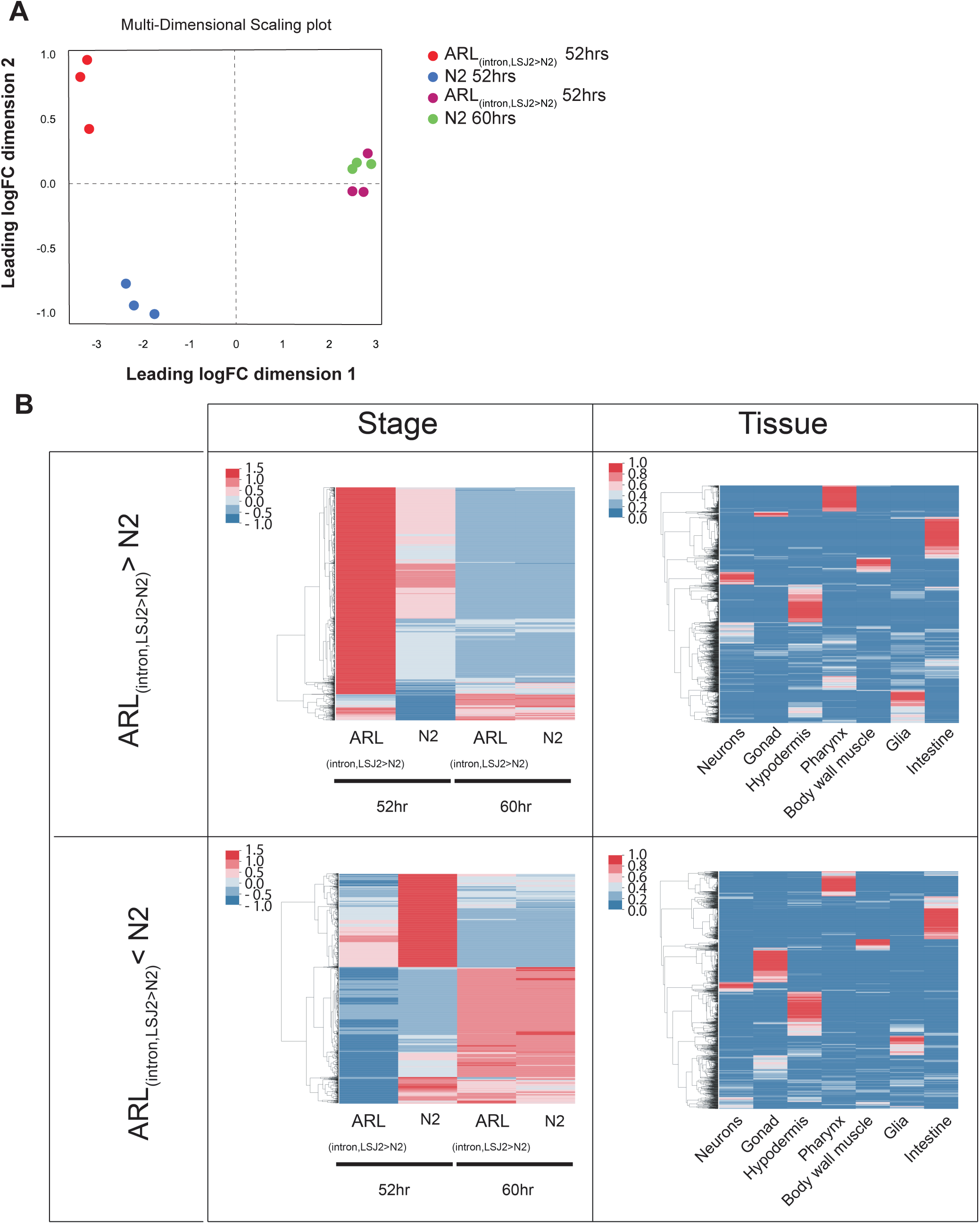
Transcriptional analysis of N2 and ARL_(intron,LSJ2>N2)_ at 52 and 60 hours. **A)** Multi-dimensional scaling analysis of transcriptional responses indicates that the N2 and ARL_(intron,LSJ2>N2)_ strains show different expression patterns at 52 hours but not 60 hours. **B)** Analysis of genes upregulated (top) or downregulated (bottom) at all four strain/conditions (left) and also by tissue-specific expression (taken from Cao et al Science (2017)).

**Figure S3.**
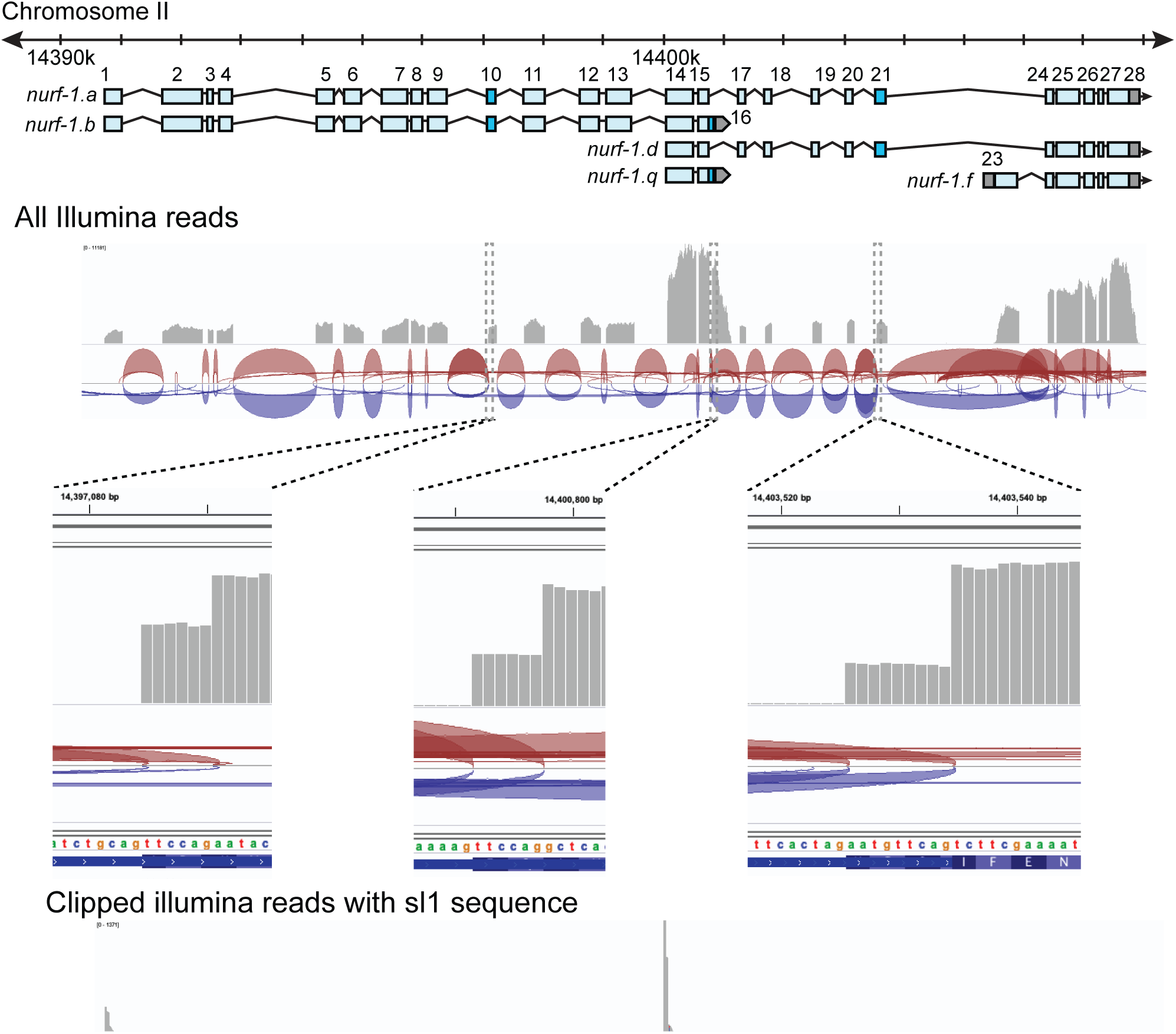
RNAseq analysis of *nurf-1*. Coverage plot of reads from RNA. Note the high expression of the 14th, 15th, and 16th exons, supporting the existance of the *nurf-1.q* transcript. Reads covering the 23rd exon were also observed, supporting the expression of the *nurf-1.f* transcript. Blow up of the 10th, 16th, and 21st exon indicates alternative splicing sites are used at these exons. Clipped reads containing sl1 sequence support transcriptional start sites at the 1st and 14th exon.

**Figure S4.**
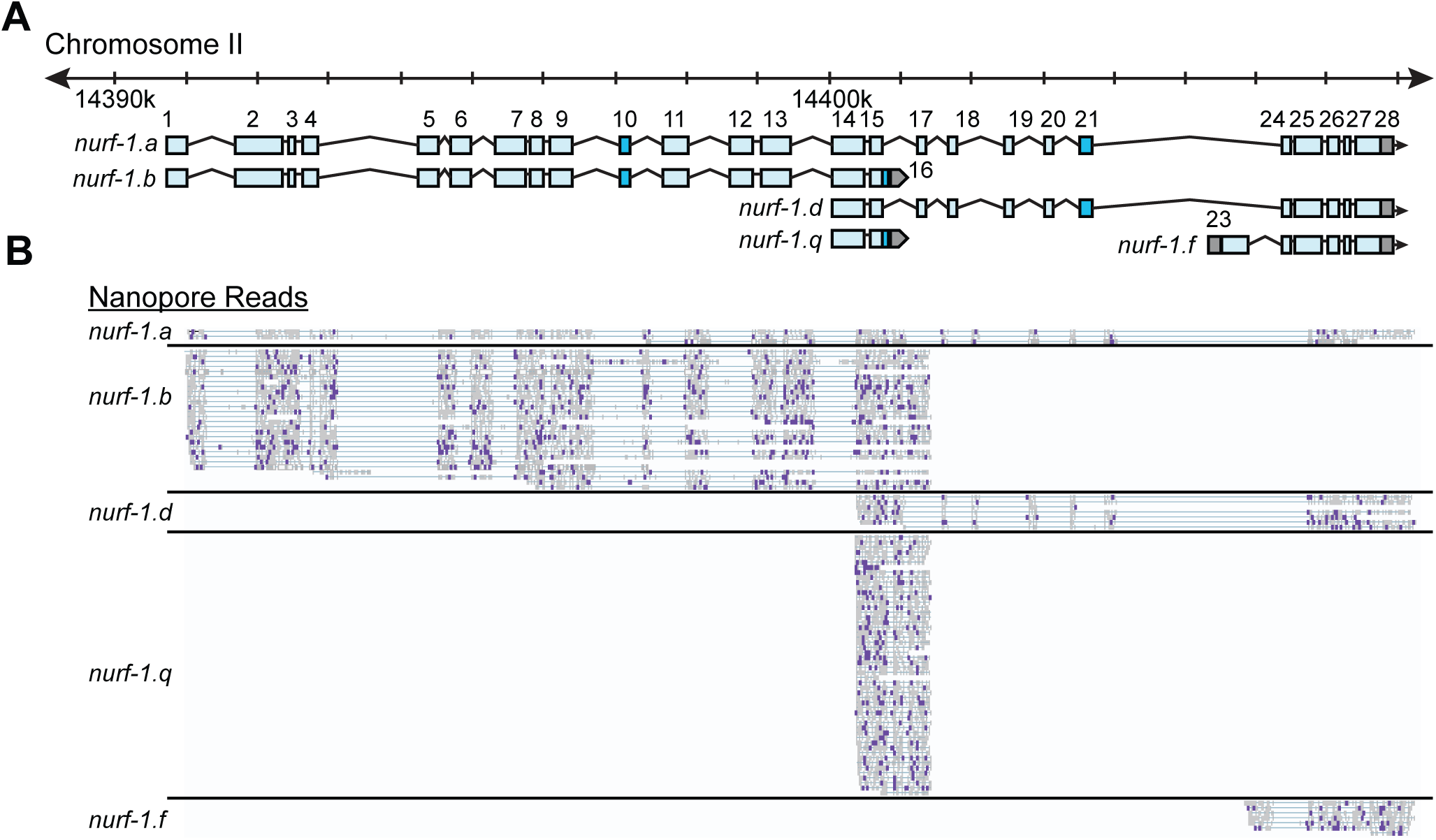
*nurf-1* encodes multiple transcripts. **A**) Subset of *nurf-1* transcripts analyzed in this paper. Each blue box is an exon. Exon number is indicated on the figure. Dark blue exons are alternatively spliced, resulting in a 6-9 bp difference in length. **B**) Nanopore sequencing reads aligned to *nurf-1*. Reads were grouped by the *nurf-1* transcripts they support. Dark purple marks are mismatches from the reference sequence.

**Figure S5.**
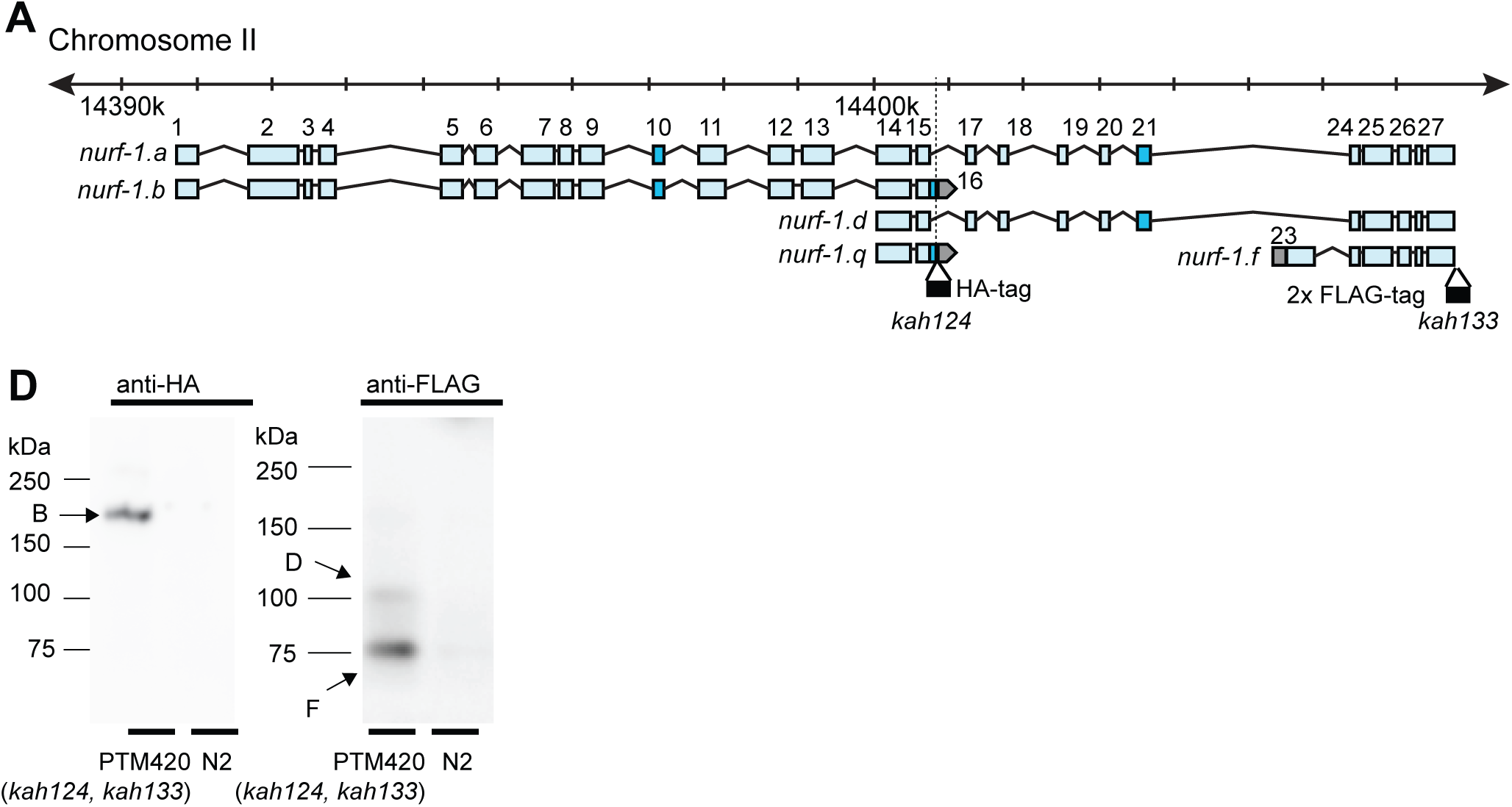
*nurf-1* encodes 3 isoforms with validated protein products. **A**) Genomic location of the HA and FLAG epitope tag insertion site are shown in black along with their associated allele names. **B**) Western blots of N2 and PTM420 strains. PTM420 contains the HA and FLAG epitope tags shown in panel A. Anti-HA antibody detected a band matching the expected size of the NURF-1.B isoform (arrow). Anti-FLAG antibody detected bands matching the expected size of the NURF-1.D and NURF-1.F isoforms (arrows).

**Figure S6.**
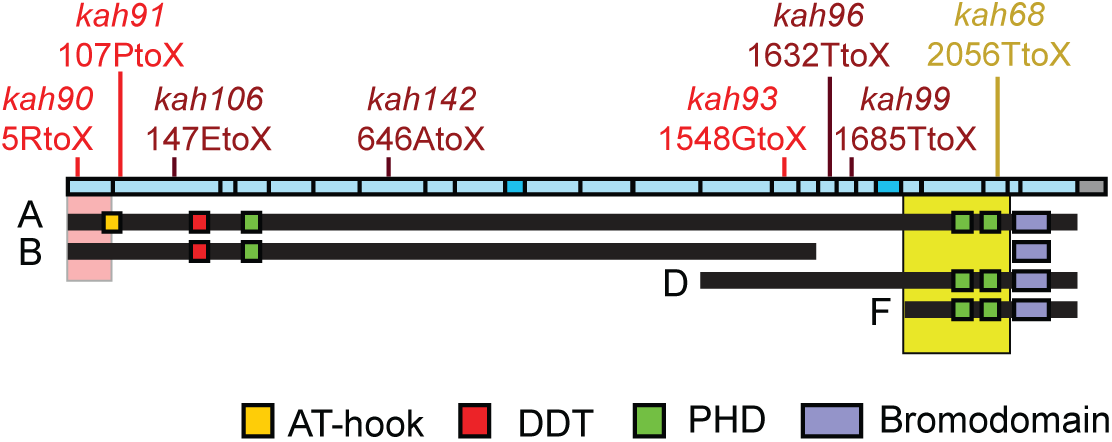
Predicted amino acid change of engineered stop codons and classical alleles on the NURF-1 isoforms.

**Figure S7.**
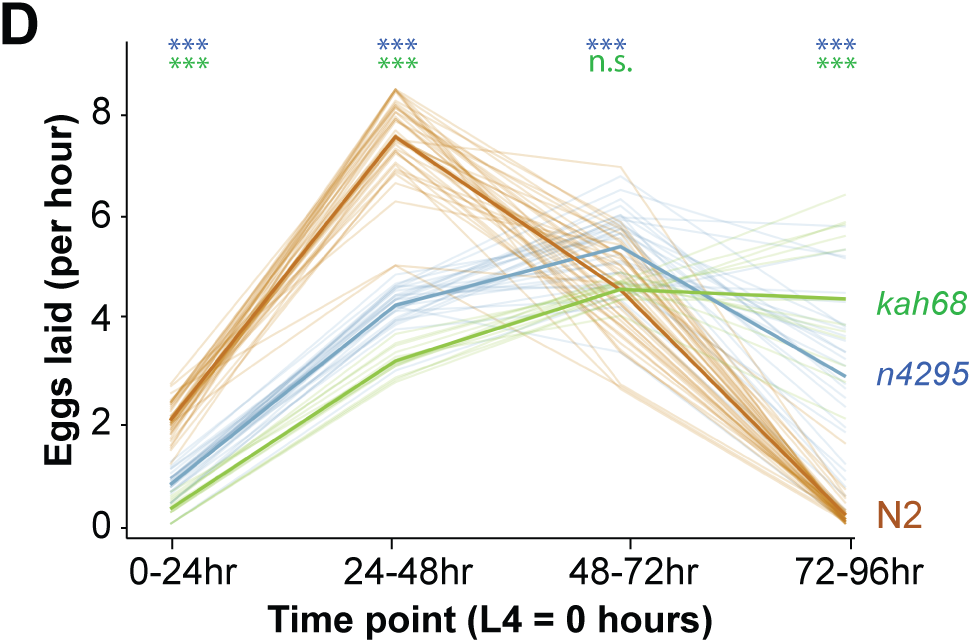
Reproductive output of indicated strains at indicated times. Six L4 animals from each strain were synchronized and placed on assay plates for the indicated timepoints. Progeny from each plate were counted to calculate the average reproductive output. Light colored lines indicate each replicate and solid colored lines show the average egg laying rate of each strain. For statistical significance, N2 vs. *kah68* comparison is in green and N2 vs. *n4295* comparison is in blue.

**Figure S8.**
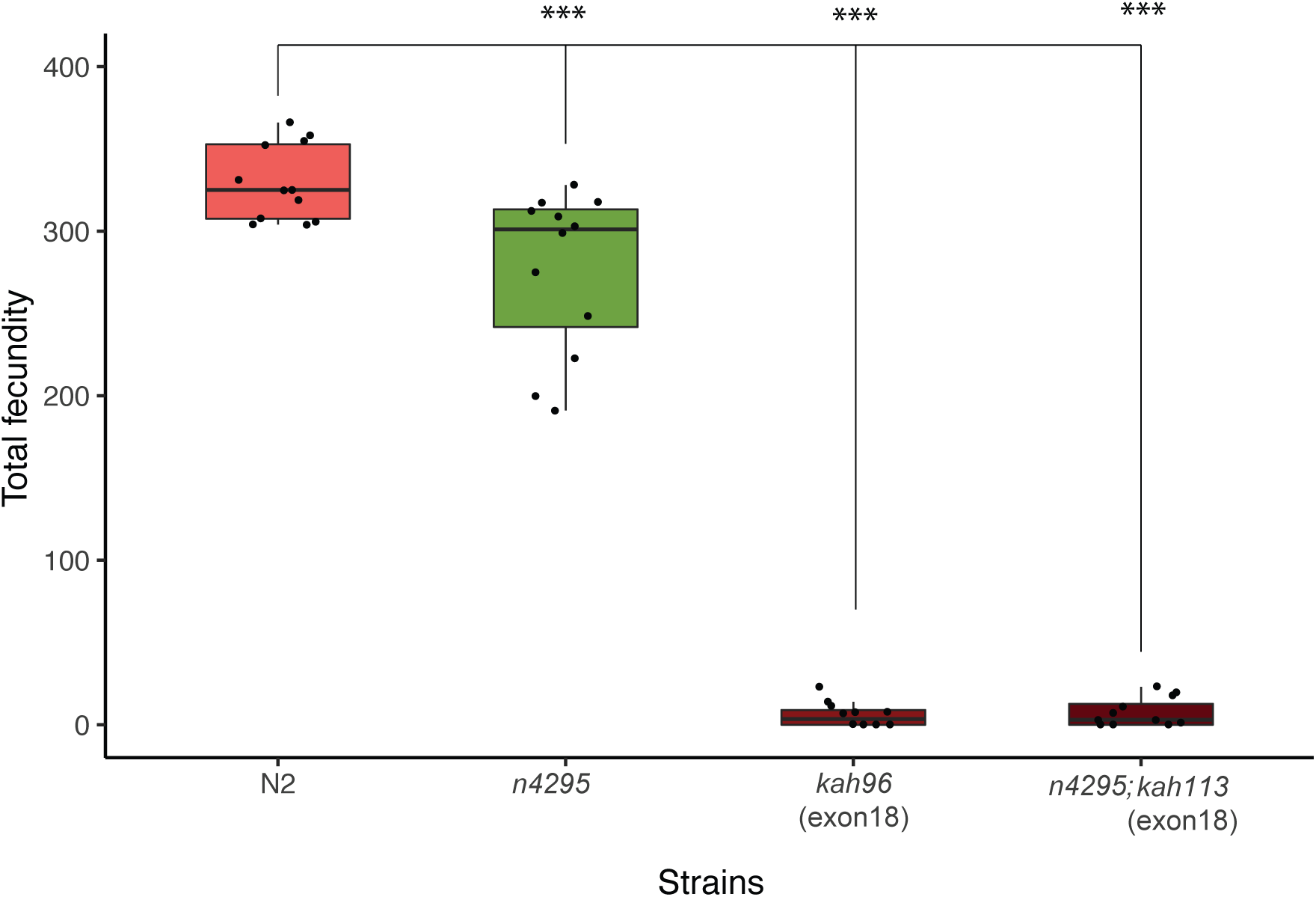
Fecundity analysis of the indicated alleles of *nurf-1*. *kah96* and *kah113* are CRISPR-induced STOP codon replacement mutations in exon 18 of *nurf-1*. Despite their identical nucleotide changes, in nomenclature, they are given unique allele names to indicate their origin from independent CRISP-Cas9 experiments.

**Figure S9.**
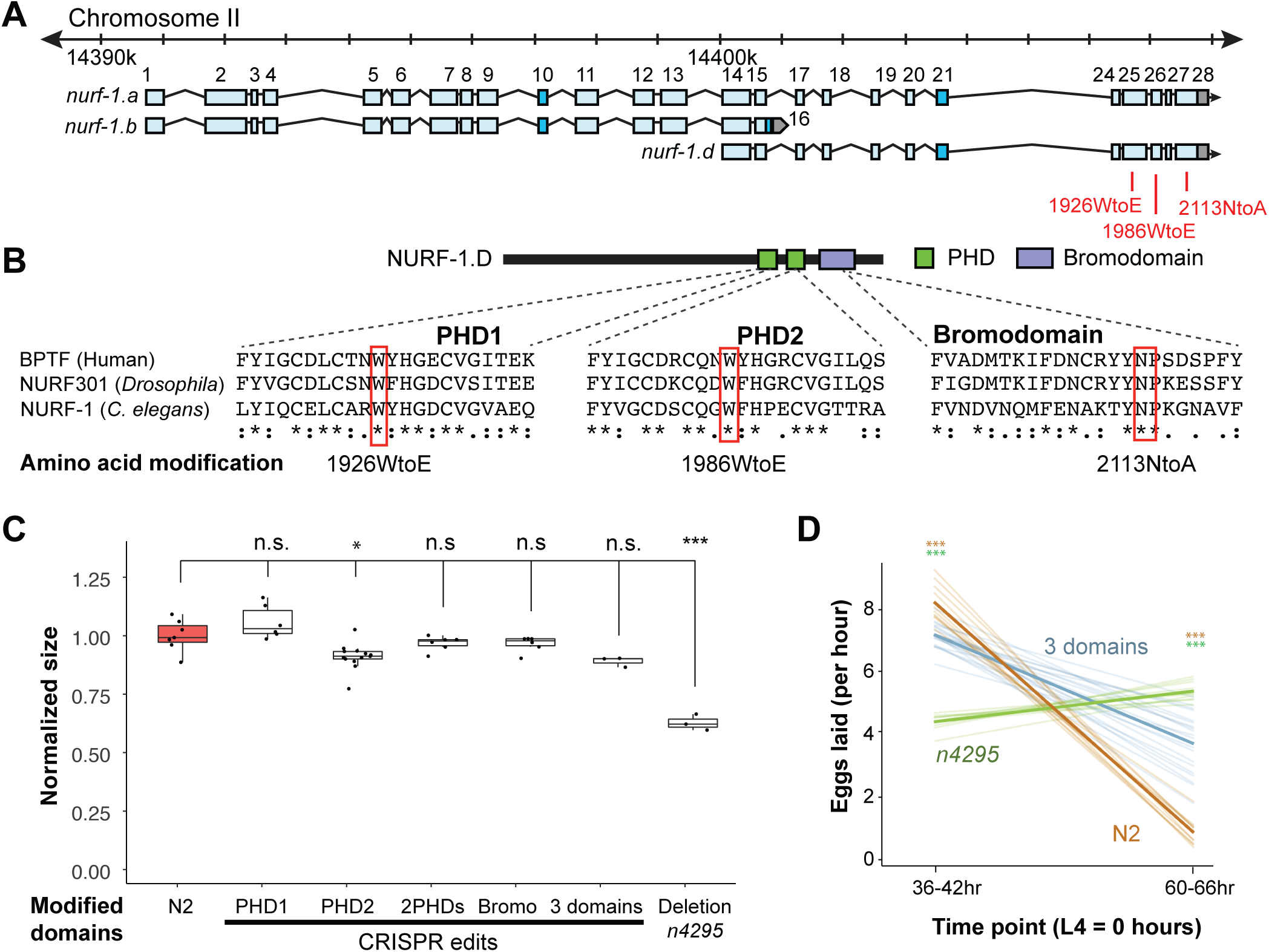
Histone recognition domains in NURF-1.D are not essential for its activity. **A**) Genomic position of CRISPR/Cas9 edits to modify conserved amino acids required for recognition of modified histone tails. **B**) Partial protein alignment of the two C-terminal PHD domains and the bromodomain showing the conserved tryptophan (W) and aspagine amino acids (N). In other species, the W to E change impairs the PHD domain’s ability to bind H3K4me3 and the N to A change impairs the bromodomains ability to bind H4K16Ac. **C**) Normalized size of indicated strains. *n4295* is a deletion allele predicted to delete both PHD domains and create a frame shift in the bromodomain. The 2PHDs strain contains both the 1,926WtoE and 1,986WtoE edits. The 3 domains strain contains all three edits shown in B. All animals were normalized to N2. **D**) Egg laying rate of indicated strains at 36-42 hours and 60-66 hours after the L4 larval stage.

**Figure S10.**
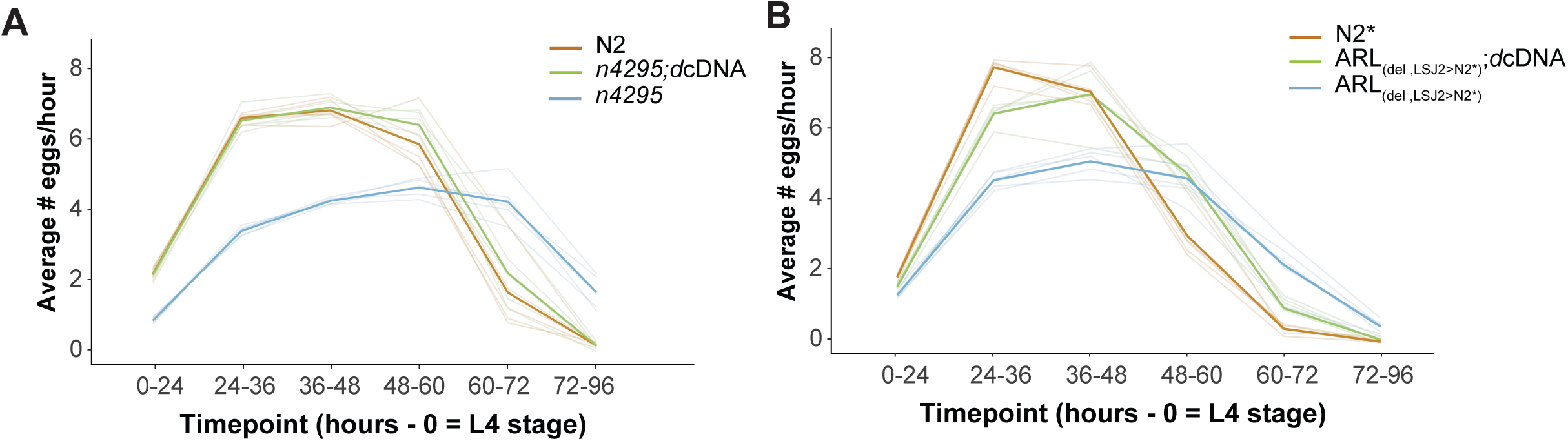
Egg-laying rate of *n4295* and ARL_(del, LSJ2>N2*)_ transgenic *nurf-1.d* cDNA rescue. Egg-laying rate was calculated at the indicated time points. Six L4 animals were picked onto each assay plates (time = 0) for the indicated times. The total number of eggs was counted for each plate and was used to calculate the average number of eggs laid per hour for each animal. Each individual trial is shown as a dim line. The mean for each strain is shown as the bold, colored lines.

**Figure S11.**
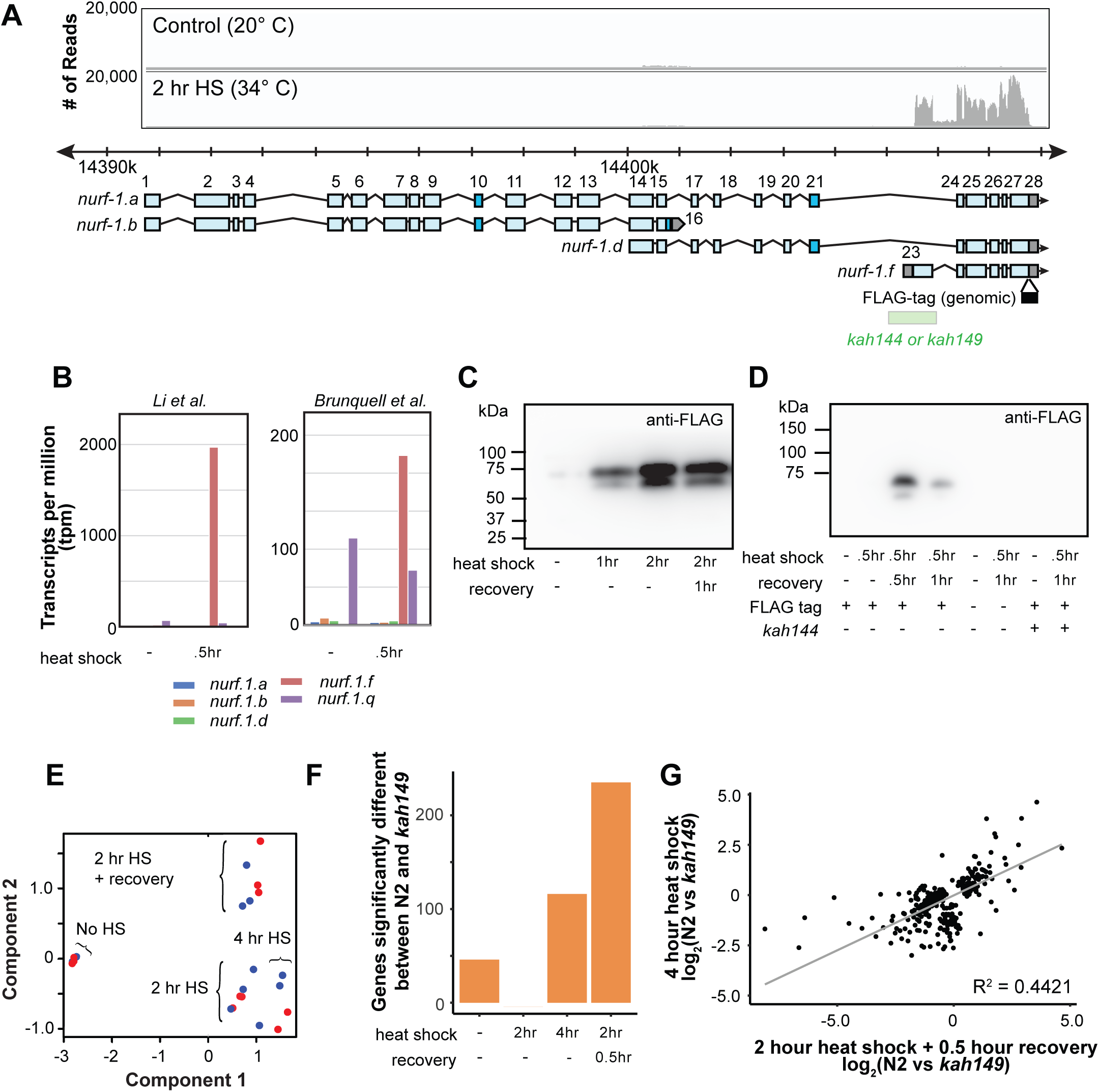
Heat shock specifically upregulates NURF-1.F. **A**) Coverage of RNAseq reads of control and heat shocked *C. elegans* animals aligned to the *nurf-1* genomic location. This data was taken from Brunquell et al. The x-axis shows the genomic location where the sequencing reads mapped, including the location of *nurf-1* transcripts. We also show the position a precise deletion of the 23rd exon edited into two strains, created using CRISPR/Cas9. In *C. elegans* genetic nomenclature, each independently generated genetic mutation is given a unique allele name, even if they are genetically identical. The deletion edited into the N2 strain is named *kah149*. The deletion edited into a strain containing an inframe FLAG epitope tag (shown as a black box) is named *kah144*. The y-axis indicates the number of reads for at each location. **B**) Quantification of RNA abundance for five *nurf-1* transcripts in response to heat shock. Data taken from Li et. al., who heat shocked L2 animals at 34°C for 30 minutes and Brunquell et. al., who heat shocked L4 animals at 33°C for 30 minutes. The y-axis is the estimated transcripts per million (tpm) for each isoform in each condition. **C**) Western blotting of a strain containing the FLAG-tag fused at the position shown in panel A using an anti-FLAG antibody. We detected two bands, one matching the predicted size of the NURF-1.F isoform, that were both upregulated by heat shock (34°C). **D**) Western blotting of three strains either containing a FLAG-tag and/or deletion allele predicted to ablate the *nurf-1.f* transcript. The x-axis shows the presence or absence of the various alleles along with the environmental condition. We detected two bands that were induced by heat shock. Observation of these bands required the FLAG epitope tag and could be ablated by the 23rd exon deletion. **E**) Multi-dimensional scaling plot (MDS) of the N2 (red) and a strain carrying the *kah149* deletion of the 23rd exon (blue) in response to various heat shock conditions. No HS indicates no heat shock, 2 hr or 4 hr HS indicates two or four hours of heat shock at 34°C, and 2 hr HS + recovery indicates animals experiencing two hours of heat shock at 34°C followed by 0.5 hours of recovery at 20°C. The overall transcriptional response was the same in both strains. **F**) Number of genes significantly up or down regulated between N2 and *kah144* animals at the indicated conditions. **G**) Scatter plot of all genes differentially expressed in four hour heat shock or two hour heat shock + recovery conditions. The R2 value was 0.4421.

**Figure S12.**
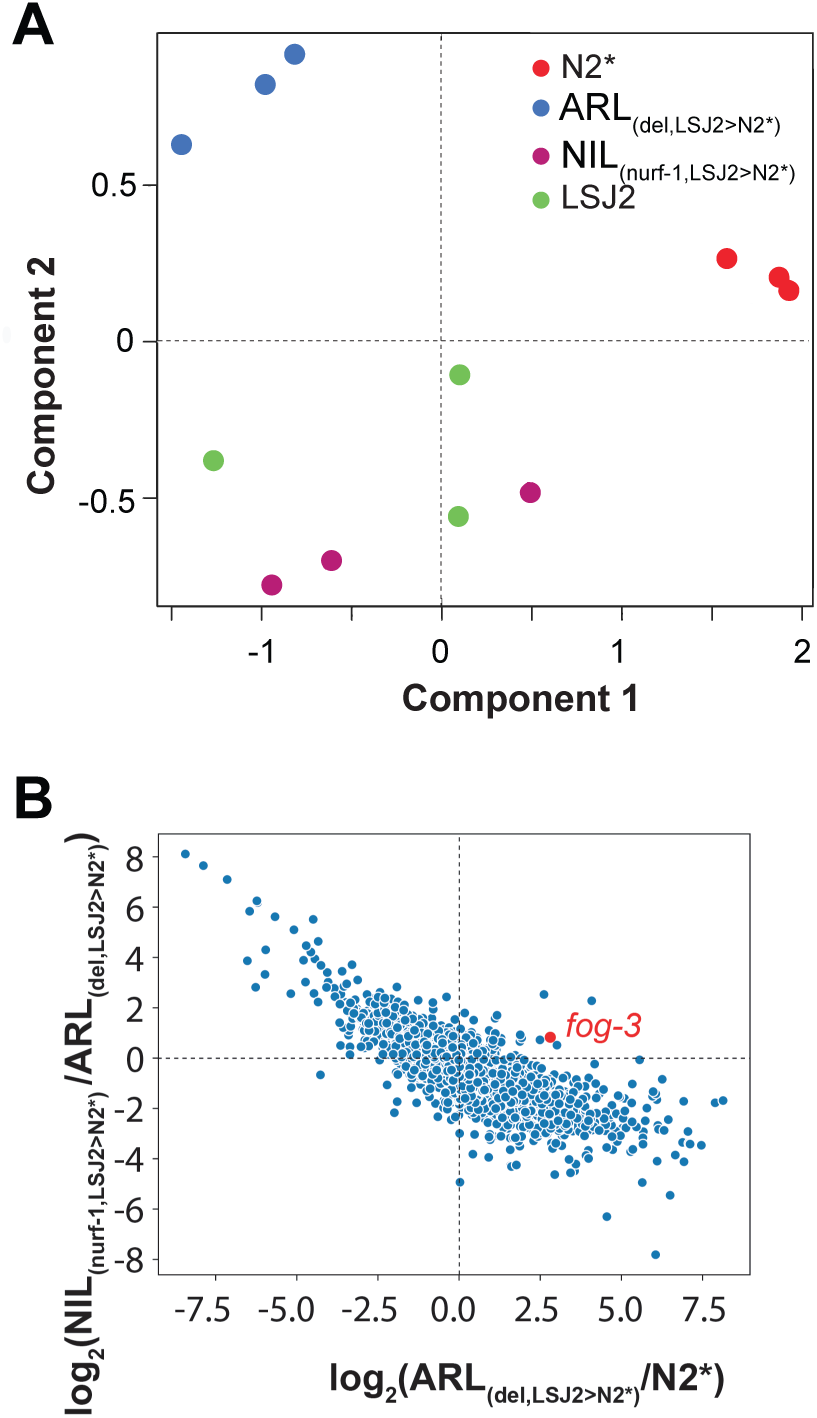
Transcriptome analysis of strains containing N2/LSJ2 genetic variation linked to nurf-1. **A**) Multi-dimensional scaling plot (MDS) of CX12311 (N2*), ARL*_del_* (PTM88), NIL*_nurf-1_* (PTM66) and LSJ2. The x-axis and y-axis are two dimensions used to separate samples from different biological conditions based upon the transcriptional change between different samples. **B**) Scatter plot of all genes detected in RNA sequencing. The x-axis is the log2 of the relative expression changes of each gene in ARLdel vs. N2* (indicating transcriptional responses induced by the 60bp deletion). The y-axis is the log2 of the relative expression of each gene in NIL*_nurf-1_* vs. ARL*_del_* (indicating transcriptional responses induced by other mutations linked to the 60bp deletion including the LSJ2-derived intron SNV). The transcriptional changes are negatively correlated with an R2 value of 0.63, indicating the genetic variation regulates a common subset of genes. *fog-3* gene is shown in red.

**Figure S13.**
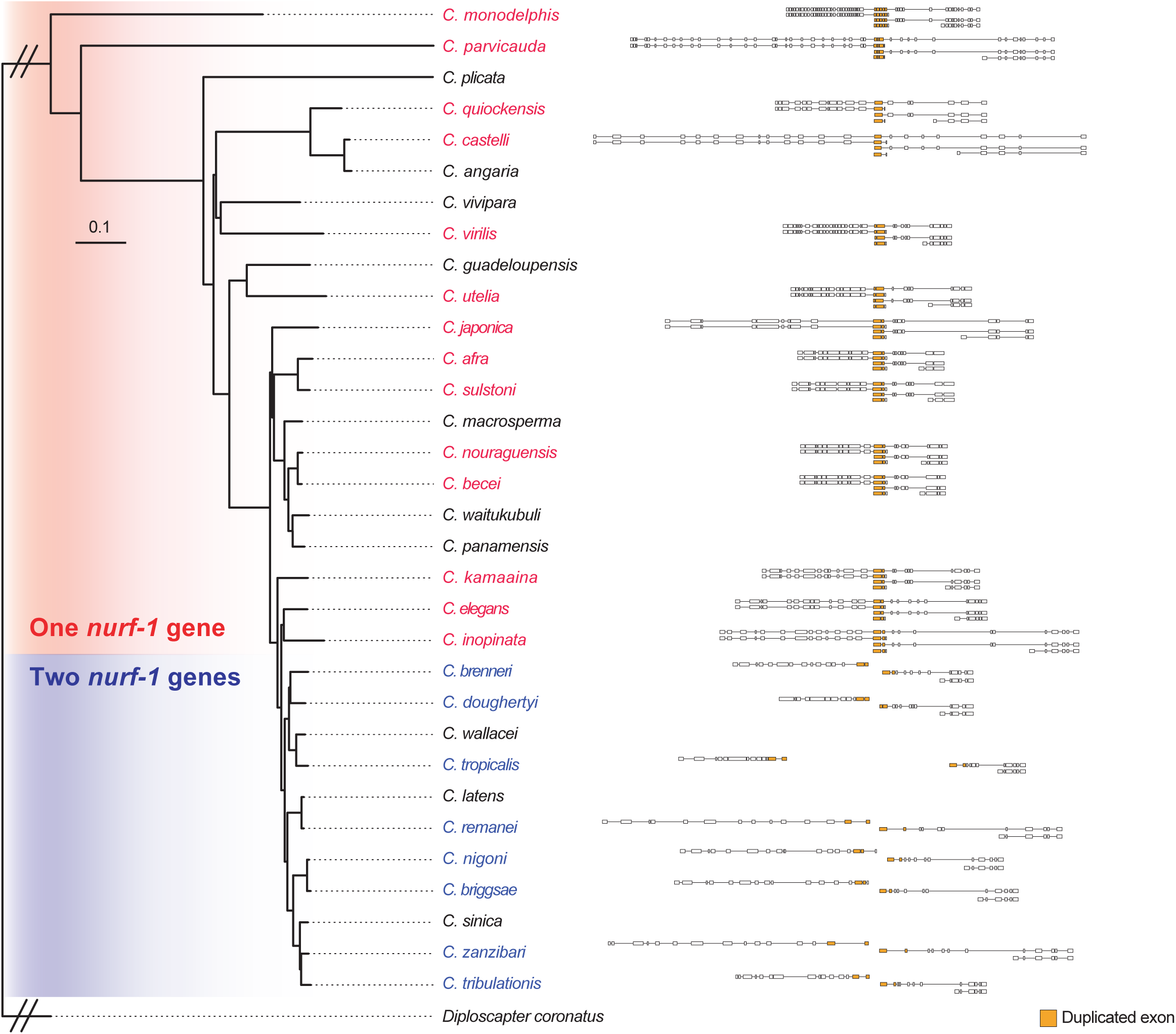
*nurf-1* isoform structure for 22 *Caenorhabditis* species. From the phylogenic tree of 32 *Caenorhabditis* species, we determined the *nurf-1* gene structure of 22 species using genome and transcriptome information. Species with one *nurf-1* gene (in red) are consistent with expression of five transcripts orthologous to *C. elegans nurf-1.a, nurf-1.b, nurf-1.q, nurf-1.d* and *nurf-1.f*. Species with two *nurf-1* genes (in blue) contain suplicated sequence and transcripts matching *C. elegans nurf-1.b (nurf-1-1), nurf-1.d (nurf-1-2.d)* and *nurf-1.f (nurf-1-2.f)*. Exons corresponding to the duplicated region were labeled in orange.

**Figure S14.**
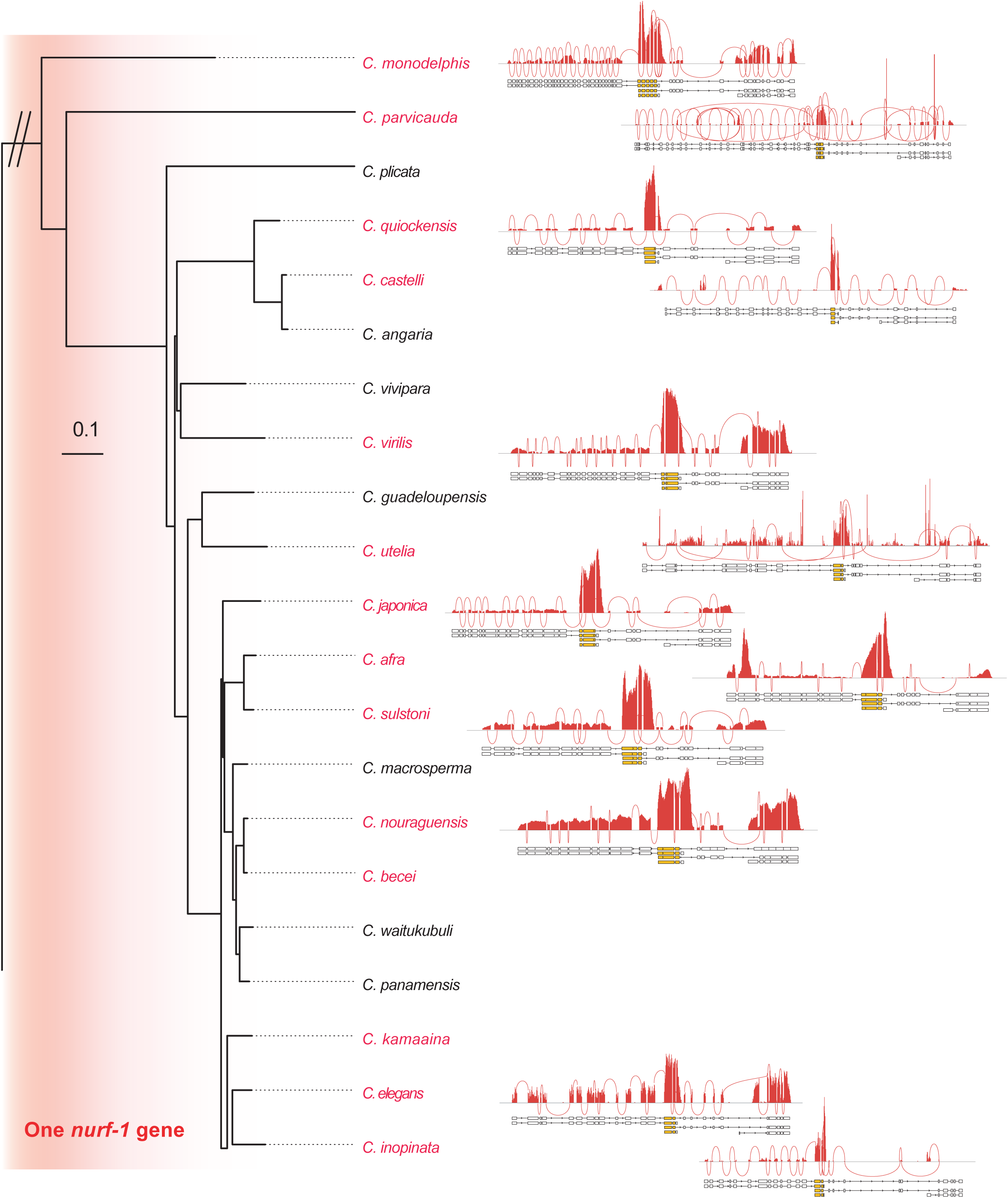
Sashimi plots for *Caenorhabditis* species with one *nurf-1* gene. Only species with published genome and transcriptome were plotted. Each peak shows the coverage for each exon, each trajectory shows exon-exon junctions supported by RNAseq reads.

**Figure S15.**
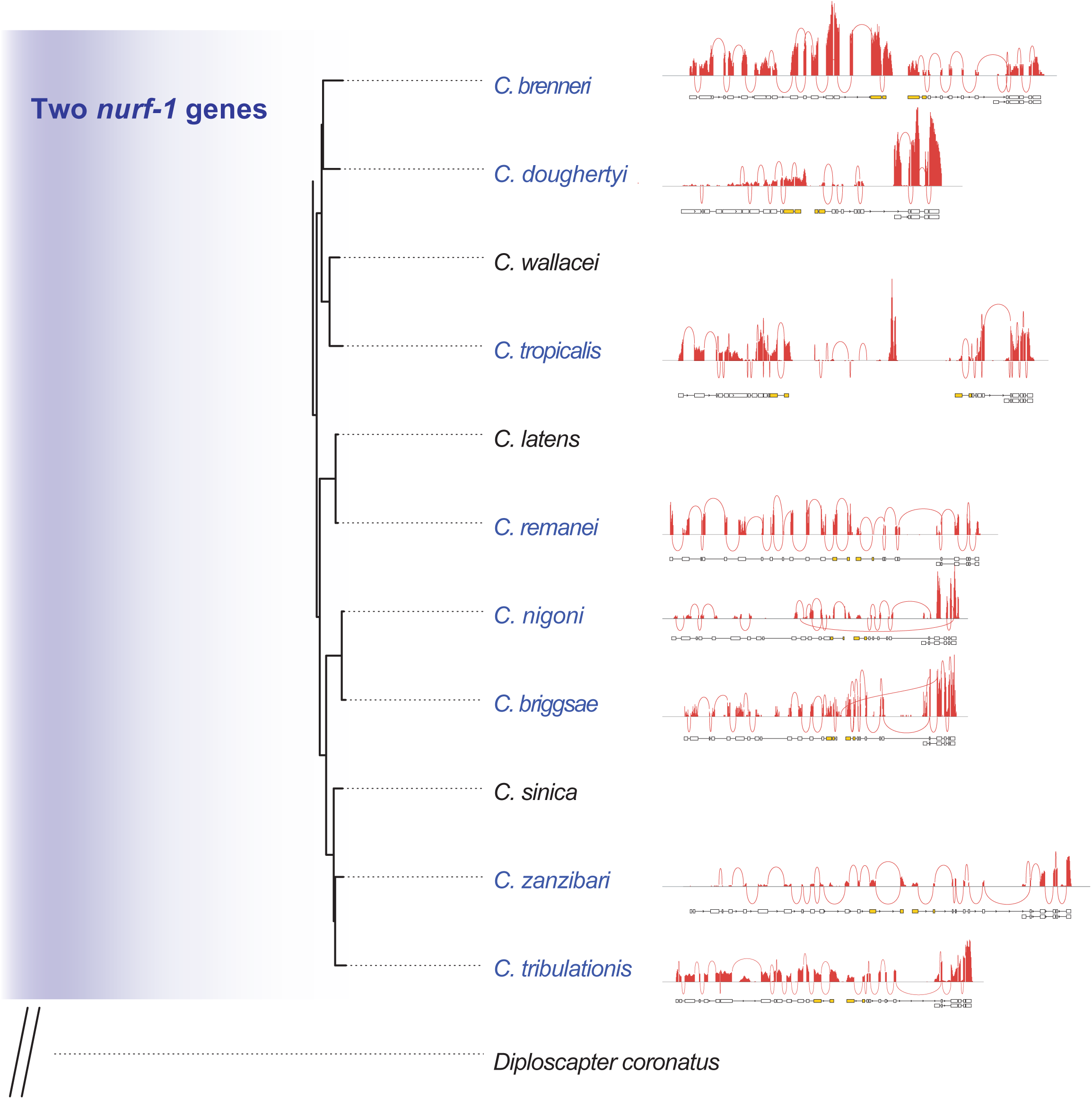
Sashimi plots for *Caenorhabditis* species with two *nurf-1* gene. Only species with published genome and transcriptome were plotted. Each peak shows the coverage for each exon, each trajectory shows exon-exon junctions supported by RNAseq reads. These plots show no read support the splicing from *nurf-1-1* to *nurf-1-2* which further suggest the split of *nurf-1* in these species.

**Figure S16.**
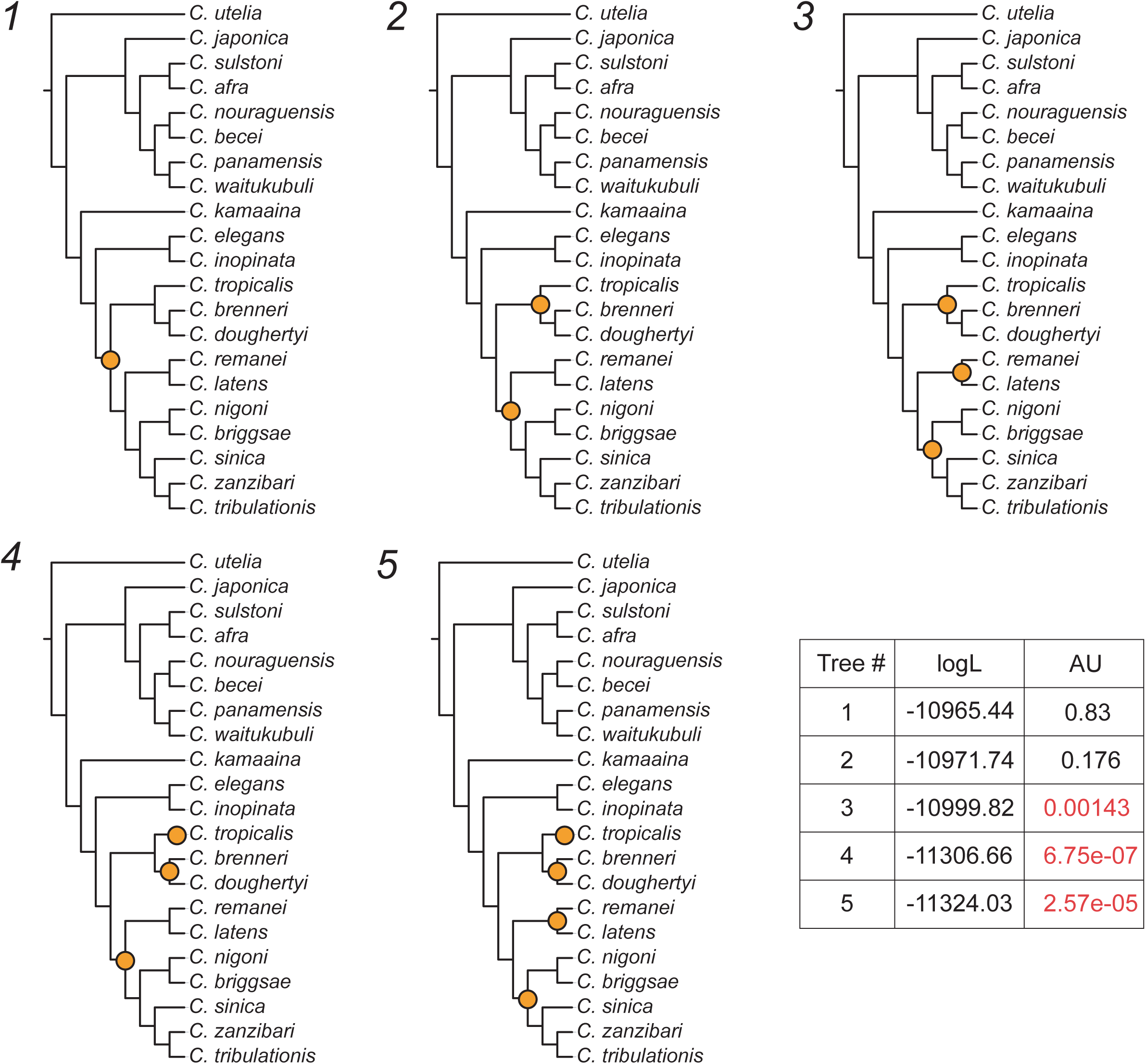
Five hypothetical topologies related to the timing and number of duplication events involved in the *nurf-1* gene split. Only those five topologies with the lowest log-likelihoods are shown. logL: log-likelihoods, AU: p-values of the approximately unbiased test. Orange circles indicate duplication events. Trees 3-5 were rejected by the AU test and are highlighted in red. Analyses were performed using IQ-TREE with the JTT substitution model with gamma-distributed rate variation among sites.

**Figure S17.**
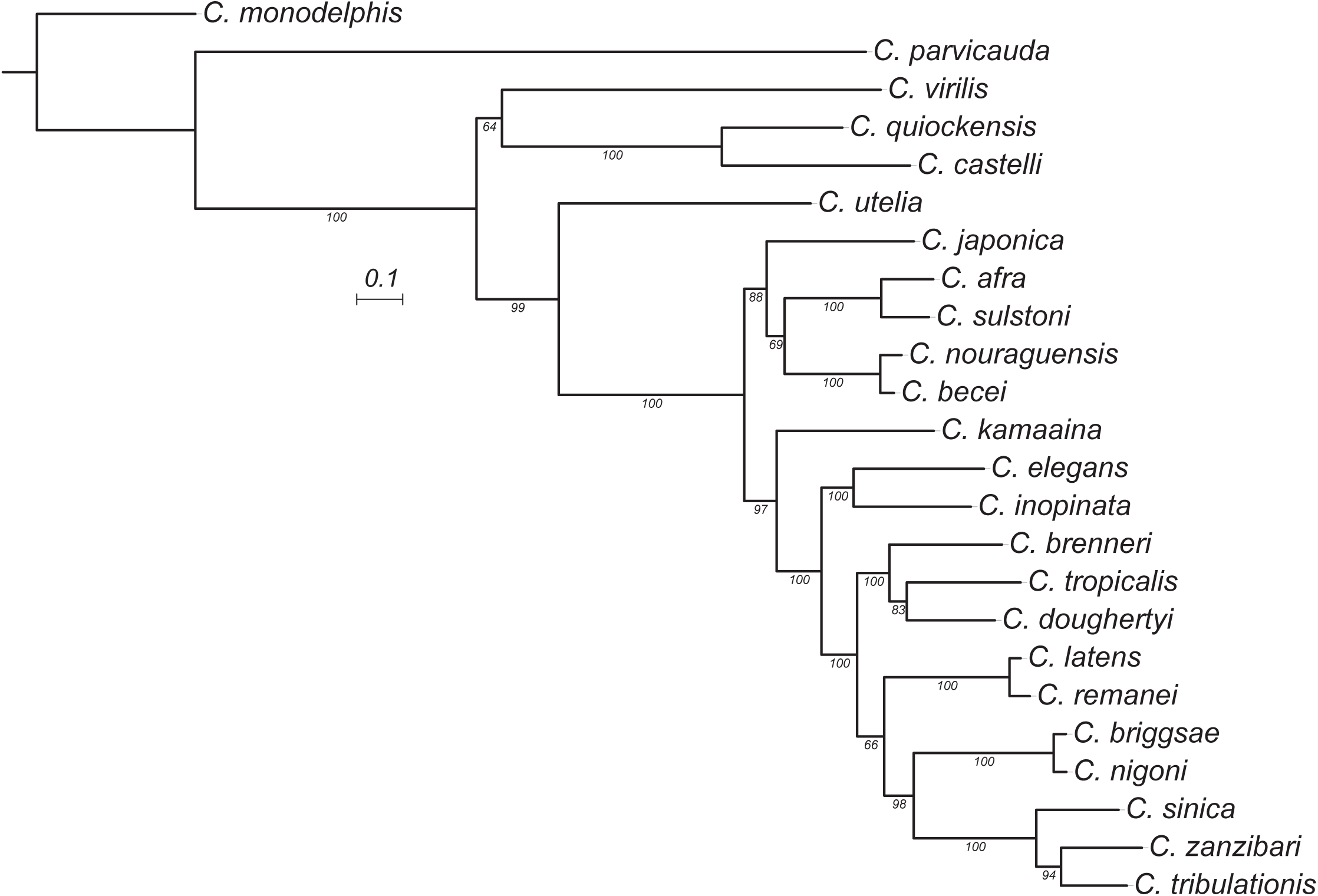
Maximum likelihood tree of the B isoform and *nurf-1-1*. The duplicated region was removed from the alignment prior to inference. Estimated using IQ-TREE using the JTT substitution model with gamma-distributed rate variation among sites. Bootstrap support values are indicated as labels on branches. Scale is in amino acid substitutions per site.

**Figure S18.**
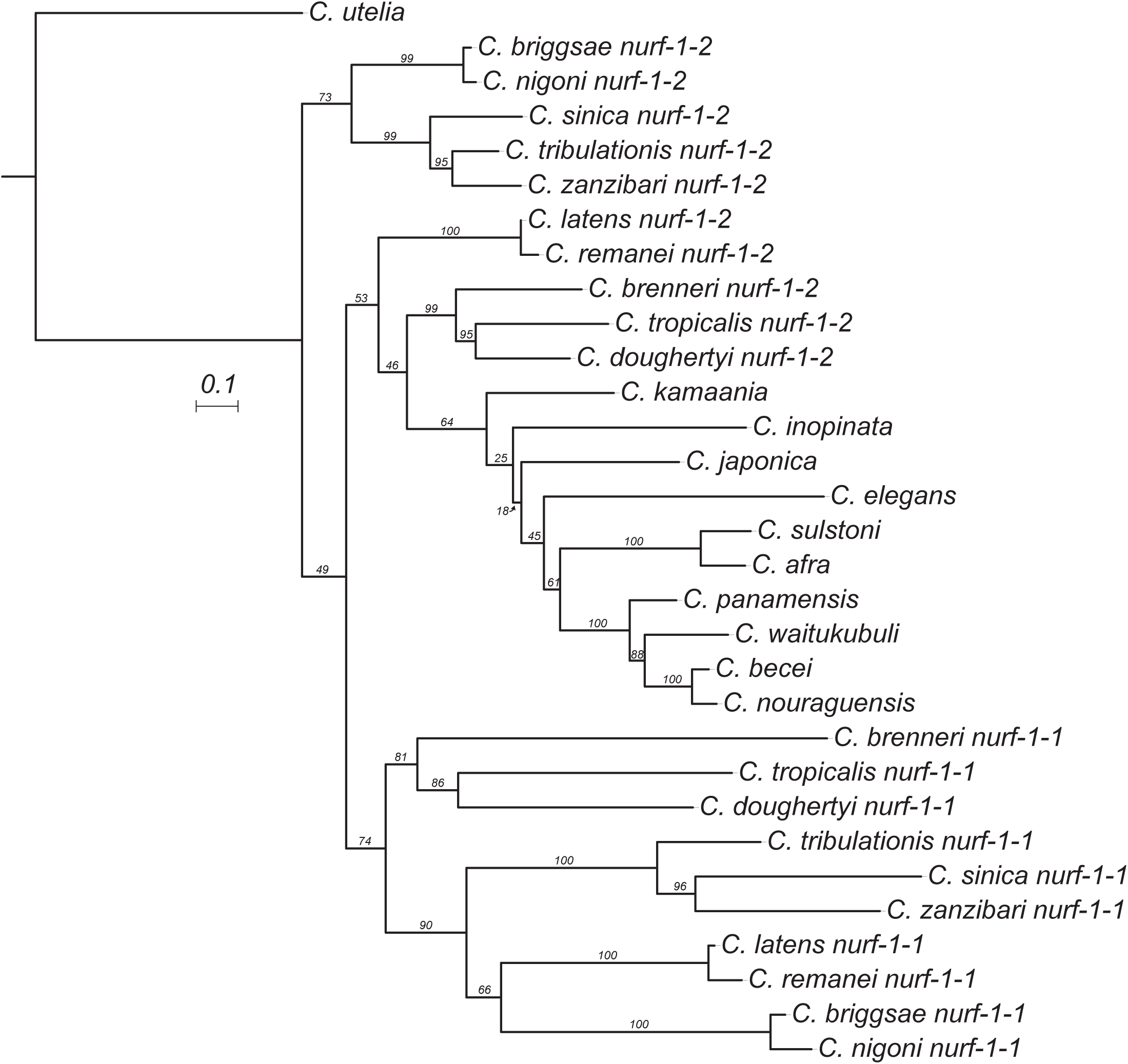
Maximum likelihood tree of the duplicated region of *nurf-1* in 22 species. Estimated using IQ-TREE using the JTT substitution model with gamma-distributed rate variation among sites. Bootstrap support values are indicated as labels on branches. Scale is in amino acid substitutions per site.

